# Trilliumosides A-B, two novel steroidal saponins isolated from Rhizomes of *Trillium govanianum* as potent anticancer agents targeting apoptosis in A-549 cancer cell line

**DOI:** 10.1101/2023.04.17.537170

**Authors:** Bashir Ahmad Lone, Misbah Tabassum, Anil Bhushan, Urvashi Dhiman, Dixhya Rani, Prem N. Gupta, D. M. Mondhe, Sumeet Gairola, Prasoon Gupta

## Abstract

Two novel steroidal saponins, Trilliumosides A (**1**) and B (**2**) were isolated from the rhizomes of *Trillium govanianum* by bioactivity-guided phytochemical investigation along with seven known compounds protodioscin (**3**) govanoside B (**4**), borassoside E (**5**), 20-hydroxyecdysone (**6**), 5-20-hydroxyecdysone (**7**), govanic acid (**8**), and diosgenin (**9**). The structure of novel compounds 1-2 were established using spectroscopic methods such as 1D, 2D NMR data and HR-ESI-MS. The isolated compounds were evaluated for *in-vitro* cytotoxic activity against a panel of human cancer cell lines. Compound **1** showed significant cytotoxic activity against A-549 (Lung) and SW-620 (Colon) cell lines with IC_50_ values of 1.83 and 1.85 µM, whereas compound (2) IC_50_ value against A-549 cell line was found to be 1.79 µM. Among previously known compounds (3), (5) and (9) their cytotoxic IC_50_ value was found to be in the range of 5-10 µM. In detailed anticancer analysis compound (2) was seen inhibiting colony forming potential and *in-vitro* migration in the A-549 cell line. Furthermore, the mechanistic study of compound (2) on the A-549 cell line revealed characteristic changes including nuclear morphology, increased ROS generation, and reduced levels of MMP. Above mentioned events eventually induce apoptosis, a key hallmark in cancer studies, by upregulating the pro-apoptotic protein BAX and downregulating the anti-apoptotic protein BCL-2 thereby activating Caspase-3. Our study reports the first mechanistic anticancer evaluation of the compounds isolated from the rhizomes of *Trillium govanianum*with remarkable activity in the desired micro molar range.

**Graphical abstract:** 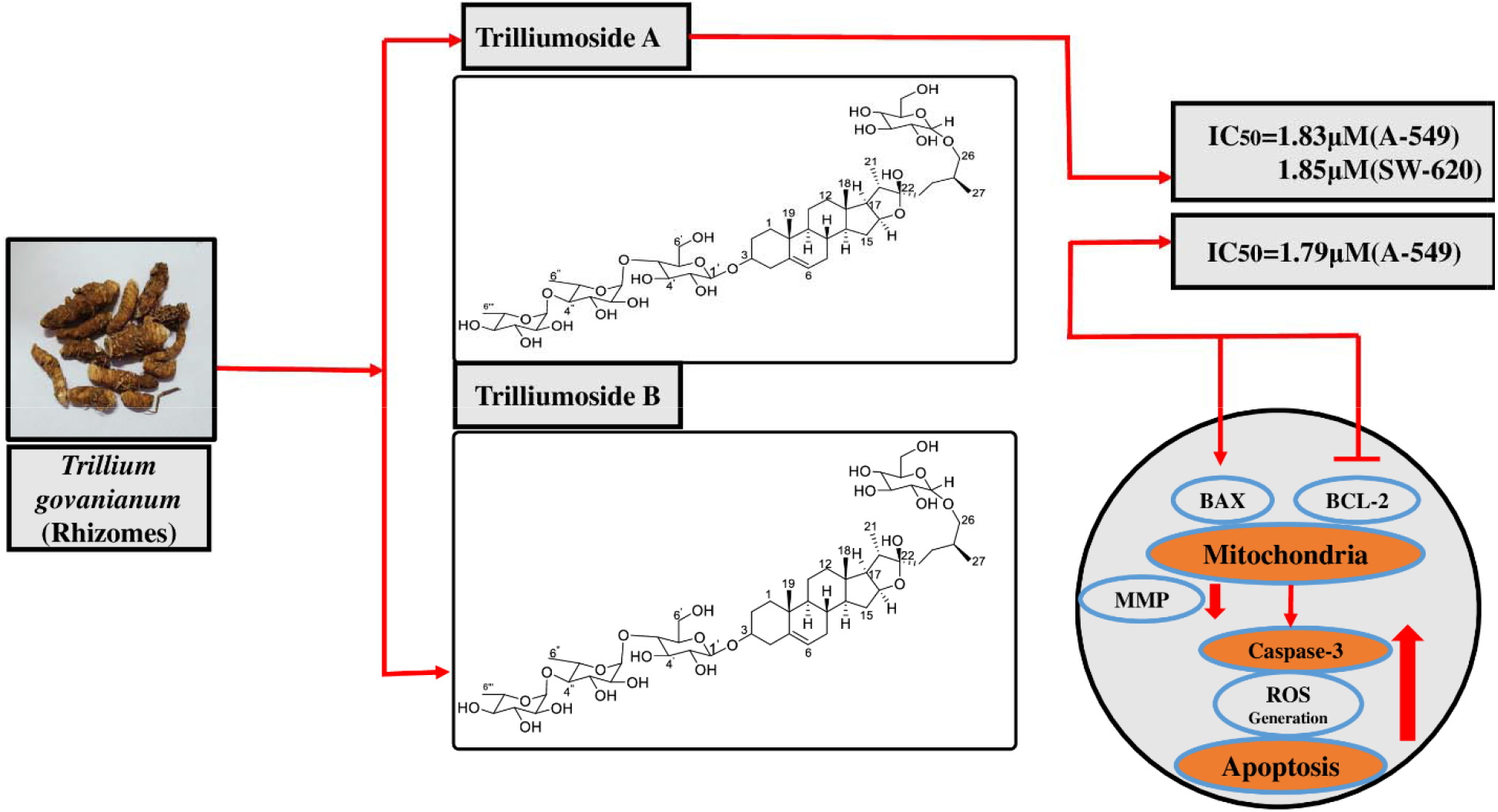

## 1. Introduction

*Trillium govanianum*(TG) is an indigenous, perennial, medicinal herb of the North-Western Himalayan region belonging to the family Melanthiaceae. Genus *Trillium* consists of forty-two species found indifferent areas across the globe. According to Global Biodiversity Information Facility (GBIF) most of the species are located in North America and Europe, and only nine are found in the Asian continent [1–2]. In India TGis commonly known by vernacular names like Nag Chhatri and distributed in an altitudinal range of 2500-3800 m in the Himalayan region [3–4].In the traditional medicine system, the rhizomes of TG are used to treatwound healing, skin diseases, dysentery, and menstrual and sexual disorders [5–6]. Increased market demand for TG at the international level is due to an essential phytosteriodsapogenin, i.e., Diosgenin (2.5%), which is an importantcomponent of commercial steroids and sex hormones [7]. Previous phytochemical investigations of genus *Trillium* have reported the identification of fatty acid esters, saponins, phenolics, terpenoids, flavonoids and steroids. Among all the isolates, steroidal saponins were found to be major bioactive compounds [8–12]. To date, more than 30 steroidal saponins have been discovered from *Trillium* species and they were found to exhibitanti-oxidant, anti-fungal, anti-inflammatory, and anti-cancer properties [13].

The steroidal saponins are important natural glycosidic compounds that possess amphiphilic character andshow remarkable cytotoxic activity [14]. The studies show that steroidal saponins exhibit anticancer properties, via its ability to decrease tumour growth, induce apoptosis, promote autophagy, and control the tumour microenvironment via several signalling pathways [15]. In our continuing efforts to discover cytotoxic compounds from plants, the crude extracts, fractions, and steroidal saponins of TG were found to have anti-cancer, anti-leishmanial, anti-bacterial, anti-fungal, anti-inflammatory and antioxidant properties [16]. The present study aimed to isolate and characterize two new steroidal saponins (1&2) and seven known compounds (3-9). Structural determination of these compounds was based on modern spectroscopic analysis, including HR-EMS, ID, 2D-NMR, and acid hydrolysis.

## 2. Materialand Methods

### 2.1 General Experimental Procedures and Chemicals Used

Column chromatography was done using silica gel (60-120 and100–200mesh),HP-20, HP-20SS, Sephedex and silver nitrate incorporated silica gel. Merck Kieselgel (Aufoilen) 60 F254 plates were used for thin-layer chromatography (TLC). Allsolvents used for HPLC analysis were bought from Merck chemicals (Mumbai).Water used for extraction and isolation purpose was obtained CentralDrug House (P) Ltd Delhi. Spectroscopic data (NMR) of isolated compounds was performed on a 400 MHz Bruker spectrometer. The reference point was TMS (*δ*_H_ and *δ*_C_: 0.00 ppm).Chemical shifts (*δ*) were referenced internally to the residual solvent peak (CD_3_OD: ^1^H-3.30, ^13^C-49.0 ppm). In 2D NMR all heteronuclear^1^H and ^13^C correlations were established on the basis of gradient-enhanced inverse-detectedHeteronuclear Multiple Bond Correlation(HMBC) and Heteronuclear single quantum coherence (HSQC) experiments. HP-20, HP-20SS, Sephedex,Silver nitrate, DMSOand all other chemicals were purchased from Sigma– Aldrich (St. Louis, MO, United States).

### 2.2 Plant Material

Rhizomes of *Trillium govanianum* were collected from Shroth Dhar, PaddarKisthwarDist. Doda of (J&K, India) between(between 3000 and 3,074 m asl).Authenticationthe crude herbwas done by Dr.SumeetGairola (Plant Science and Agrotechnology DivisionIIIM), the voucher specimen (No. RRLH-23418) was submitted in the Crude Drug Repository of CSIR-IIIM, Jammu.

### 2.3 Extraction and isolation

Shade dried rhizomesof*Trillium govanianum* (1.9 kg) were grinded into powder and sequentiallyextracted twice (drenched for 12 h each), with 3.5 L of chloroform followed by 3 L 20:80 (H2O: MeOH) thrice. The extractswere filteredseparatelyandconcentrated using rotavaporat 45^◦^Cat reduced pressure to obtain a crude chloroform and hydroalcoholicextract (187.78 and 229.44g), respectively. Further (20%aq. MeOH) extract (229.44 g) was subjected to reverse phase column chromatography for purification using HP-20 resin eluted with gradient of water: methanol solvent system (100:00.0–10:90.0 one liter collected measurements of all fractions) and on concentration, giving 20 collective subfractions (HA-01-HA-20). Further, the white precipitate was formed in Fr. HA-12-HA-16), which on separation with Whatman filter paper yielded white amorphous powder **compound 5** (8.67 g), which was detected on TLC [CHCl_3_: MeOH: H_2_O (6.5:3:0.5)] as a single dark green spot onheatingthe dried TLC in the anisaldehyde sulphuric acid reagent. Purification of sub fraction HA-01-HA07(16.45 g) on (HP-20SS) resin, using a gradient of MeOH: H_2_O (95 to 55%), afforded 35 small fractions (100 ml each), and these fractions were divided into six subfractions on the bases of their chemical profiling namely (F-1A to F-1F).Fraction F-1D (1.2 g) on purification using HP-20SS eluted with water: methanol solvent system (0.5:9.5–8.5:2.525 ml collected in 50 mL tubes), based on the TLC profile fractions (6.5:3.5), (8.5:2.5) afforded **compound 1 and 2** (27, and 17mg) as a white amorphous powder and dark brown crystalline solid respectively. Also, purification of fraction F-1A-1B (0.9 g) on HP-20SS, using H_2_O: MeOH solvent system (70:30), afforded **compound 3** (40 mg) as white solid powder, remixing (95:0.5-75:2.5) fractions (291 mg) yielded **compound 4** (61 mg) yellowish powder. Chloroform extract (187.78) was processed for fractionation to separate oily non polar part using solvent–solvent fractionation. Extract (187.78 gm) was dissolved in 1 Litre of CHCl_3_ and 800 mL of water was added to solubilise the extract,equal amount of cyclohexane was used for, partition and hexane layer was separated, concentrated using rotavapor (20.1 gm) oily mass was left behind. The hexane fraction/extract was processed for purification by CC (Silver nitrate incorporated silica gel, 100–200 mesh) Hexane: Ethyl acetat egradient solvent system was used to elute (100:0 to 70:30, 200 mL fractions were collected), concentrated using rotavapor giving a total of 11 pooled fractions (F.H1– F.H11) based on their TLC outline, fraction (F.H6-H9) gives a single spot after charring and drying in the anisaldehyde reagent **compound 6** (White powder1.66 gm). Remaining chloroform residual extract(131.66 gm) was processed for isolation of individual compounds by CC (silica gel, 100–200 mesh), eluted with an increasing polarity of Hexane:Ethyl acetate (500 mL collected volumes). Eluted fractions were concentrated on rotavapor giving six new fractions (F.CR-1– F.CR-6) based on the TLC analysis. Fraction CR-03 (17.78 g) was again processed for purificationby CC using(silica gel 100-200 mesh) with n-hexane-ethyl acetate (1:0 to 8:2, 100 mLvolumes were collected) toobtain nine daughter fractions (3A-3I). On placing these fractions overnighton working table,a colourless needleswere formed in Fr. 3D-3G yielded **compound 7**(1.11g), which was visualisedusing developing solvent system of(Hex-EtOAc 80:20), a bright green spotappearedafter sprayinganisaldehyde reagentand heatingTLC plate. Moreover CR-4-6 (28.45 g)fraction was again purifiedby CC on silica gel (60–120 mesh, 200 g), using a gradient of CHCl3–MeOH (100 to 80%), a total of 22 fractions were obtained (100 ml each). These fractions were separated into six subfractions (F.A1-F6) based on similarities in their TLC profiles. The several times repeated CC of F.B2 fraction (7.27 g) on silica (230–400 mesh, 100 g), using increasing polarity of CHCl3–MeOH (85:15), afforded **compounds 8 and 9** (0.9 gm) and (42 mg), respectively. Chemical structures of all the isolated and characterized compounds are presented in figure 2.

**Compound (1):** was isolated as a brownish solid (27 mg). Its molecular formula was determined to be C_51_H_85_O_23_, by LC/MS data at (m/z 1064.13 [M+H]^+^ (calcd for C_51_H_85_O_23_^+^,1064.1911), together with its NMR data (Tables 1 and 2).

**Table 1:** 1H (500 MHz) ^13^C (100 MHz) NMR data for aglycones of 1-2 in CD_3_OD.

|  | **1** |  | **2** |  |
| --- | --- | --- | --- | --- |
| Position | $\delta_H$ (int., mult., *J* in Hz) | $\delta_C$ | $\delta_H$ (int., mult., *J* in Hz) | $\delta_C$ |
| 1a | 1.09 (1H, *m*) | 38.67 | 1.06 (1H, *m*) | 38.68 |
| B | 1.82 (1H, *m*) |  | 1.87 (1H, *m*) |  |
| 2a | 1.91 (1H, *m*) | 30.55 | 1.90 (1H, *m*) | 31.48 |
| B | 1.62 (1H, *m*) |  | 1.65 (1H, *m*) |  |
| 3 | 3.53 (1H, *m*) | 79.45 | 3.39 (1H, *m*) | 79.93 |
| 4a | 2.29 (1H, *m*) | 39.58 | 2.43 (1H, *m*) | 39.60 |
| B | 2.44 (*m*, 1H) |  | 2.27 (1H, *m*) |  |
| 5 |  | 141.95 |  | 142.00 |
| 6 | 5.38 (1H, *br, d*) | 122.75 | 5.38 (1H, *br, d*) | 122.75 |
| 7a | 1.98 (1H, *m*) | 32.33 | 2.00 (1H, *m*) | 33.29 |
| B | 1.35 (1H, *m*) |  | 1.53 (1H, *m*) |  |
| 8 | 1.54 (1H, *m*) | 33.18 | 1.63 (1H, *m*) | 32.88 |
| 9 | 0.94 (*m*, 1H) | 51.52 | 0.91 (1H, *m*) | 51.87 |
| 10 |  | 38.01 |  | 38.15 |
| 11a | 1.55 (1H, *m*) | 21.18 | 1.55 (1H, *m*) | 22.07 |
| B | 1.51 (1H, *m*) |  | 1.54 (1H, *m*) |  |
| 12a | 1.91 (1H, *m*) | 30.55 | 1.57 (1H, *m*) | 30.86 |
| B | 1.23 (1H, *m*) |  | 1.91 (1H, *m*) |  |
| 13 |  | 46.12 |  | 46.61 |
| 14 | 1.68 (1H, *m*) | 53.92 | 1.12 (1H, *m*) | 57.87 |
| 15a | 1.55 (1H, *m*) | 33.01 | 1.57 (1H, *m*) | 29.07 |
| B | 1.98 (1H, *m*) |  | 1.98 (1H, *m*) |  |
| 16 | 4.11 (1H, *dd*, 6.1, 3.2) | 90.89 | 4.36 (1H, *dd*, J, 6.1, 3.2) | 82.53 |
| 17 |  | 91.62 | 1.72 (1H, *m*) | 65.14 |
| 18 | 0.86 (3H, *s*) | 17.50 | 0.82 (3H, *s*) | 20.01 |
| 19 | 1.36 (3H, *s*) | 20.10 | 1.04 (3H, *s*) | 17.44 |
| 20 | 2.38 (1H, *m*) | 50.00 | 2.15 (1H, *m*) | 41.93 |
| 21 | 0.92 (3H, *s*) | 9.87 | 0.83 (3H, *s*) | 16.97 |
| 22 |  | 112.46 |  | 114.10 (□ - OH) |
| 23a | 1.69 (1H, *m*) | 36.88 | 1.73 (1H, *m*) | 40.94 |
| B | 1.67 (1H, *m*) |  | 1.83 (1H, *m*) |  |
| 24a | 1.18 (1H, *m*) | 28.55 | 1.14 (1H, *m*) | 28.75 |
| B | 1.68 (1H, *m*) |  | 1.60 (1H, *m*) |  |
| 25 | 1.73 (1H, *m*) | 35.07 | 1.73 (1H, *m*) | 35.08 |
| 26a | 3.41 (1H, *m*) | 58.43 | 3.38 (1H, *m*) | 76.12 |
| B | 3.72 (1H, *m*) |  | 3.70 (1H, *m*) |  |
| 27a | 3.37 (1H, *m*) | 76.11 | 1.73 (3H, *m*) | 76.11 |
| B | 3.70 (1H, *m*) |  |  |  |

**Table 2:**
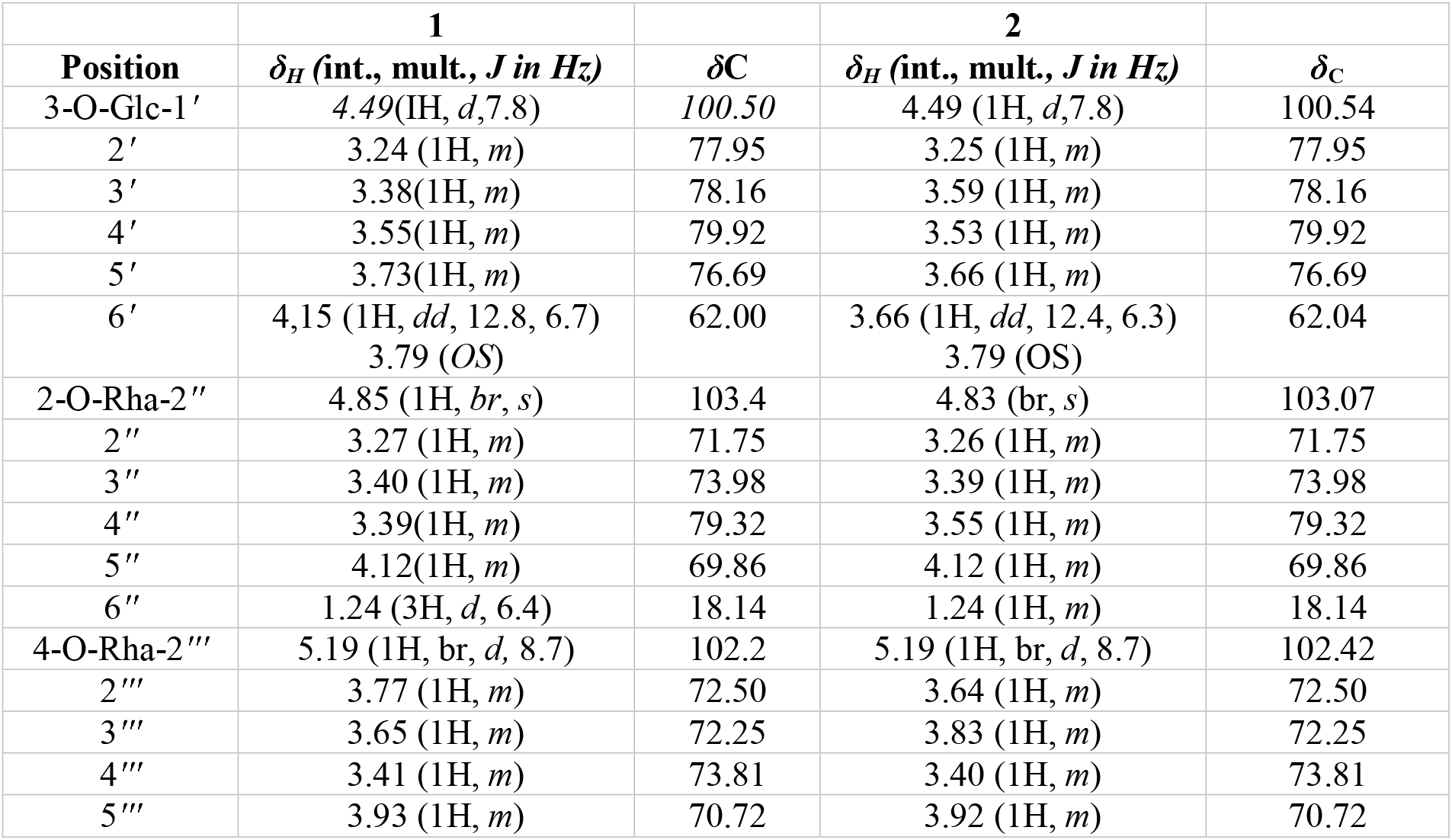
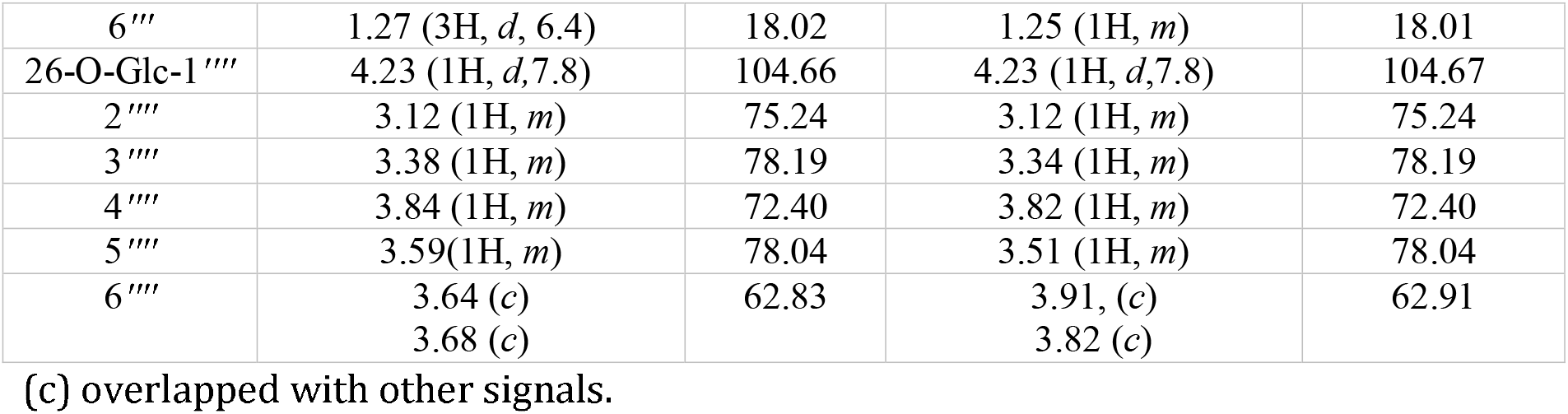
1H (500 MHz) ^13^C (100 MHz) NMR data for glycones of 1-2 in MeOD.

**Compound (2):** was isolated as a brownish amorphous solid (17 mg). Its molecular formula was determined to be C_51_H_85_O_22_, by HR-ESI-MS data at (m/z 1049.5532 [M+H]^+^ (calcd for C_51_H_85_O_22_^+^, 1049.5411), together with its NMR data (Tables 1 and 2).

**Govanoside B (3):** White amorphous powder, (37 mg);HR-ESI-MS (Positive): 1240.51 (M+Na)^+^(Calcd. C_55_H_86_O_28_Na^+^, 1240.5174);^1^H NMR (400 MHz, CD_3_OD) and ^13^C NMR (100 MHz, CD_3_OD) observed data was quite matching to that of reported in the literature and compound was identified as BorassideE[17].

**Protodioscin (4):** White colour amorphous solid, (40 mg); ESI-MS: (Positive) *m/z* 1071.43 [M+Na]^+^(Calcd. for C_51_H_84_O_22_Na^+^, 1071.43); ^1^H-NMR (CD_3_OD, 400 MHz) and ^13^C-NMR (CD_3_OD, 100 MHz). Observed data were compared with the literature, and the compound was identified as Protodioscin [18].

**Borassoside E (5):** White colour amorphous solid, (8.67 gm); ESI-MS (Negative): 867.46 (M-H)^-^ (Calcd. 867.48, C_45_H_71_O_16_^-^);^1^H NMR (400 MHz, CD_3_OD) and ^13^C NMR (100 MHz, CD_3_OD) observed data was quite matching to that of reported in literature and compound was identified as Borassoside E [16 and19].

**Govanic acid (6):**White solid,(1.11 gm) ESI-MS(Positive); 331.2436 (M+H)^+^(calcd., C_18_H_35_O_5_^+^, 331.2436)^1^H NMR (400 MHz, CDCl3), and ^13^C NMR (100 MHz, CDCl3) observed data was matched to that of reported data, and the compound was identified as Govanic acid [11].

**Diosgenin (7):** Colourless crystalline, solid (678 mg); ESI-MS (Positive): 415 (M+H)^+^(Calcd. 415.31, C_27_H_43_O_3_^+^);^1^H NMR (400 MHz, CDCl_3_) and ^13^C NMR (100 MHz, CDCl3) observed data was similar to that of reported in the literature and compound was identified as Diosgenin [16].

**20-Hydroxy ecdysone (8):** White colour solid, (0.9g)HR-MS *m/z* 481.3161 [M+H] ^+^(calcd for C_27_H_45_O_7_^+^, 481.3161).^1^H NMR (400 MHz, CD_3_OD) and ^13^C NMR (100 MHz, CD_3_OD) observed data was matched to that of reported data and compound was identified as 20-Hydroxy ecdysone [11].

**5,20-Hydroxy ecdysone(9):** White colour solid, (40 mg)ESI-MS(Positive); 497.30 (M+H)^+^(calcd. C_18_H_45_O_8_ ^+^, 497.31, ^1^H NMR (400 MHz, CDCl_3_), and ^13^C NMR (100 MHz, CDCl3) observed data was matched to that of reported data, and the compound was identified as 5,20-Hydroxy ecdysone[11].

### 2.4 Sugar analysis of compounds (1-2)

#### 2.4.1 Acid hydrolysis and GC/MS analysis

Both new Compounds **1**and **2**(10 mg) each were dissolved in 5 mL MeOH, and then 5 mL 5% HCl (V/V) was added to the solution. The reaction mixture was refluxed on oil bath at 100°C for 3 hours after the completion of the reaction. To separate the aglycone component, methanol was distilled off, and the reaction mixture was agitated with CHCl_3_. Using rotavapor, the CHCl_3_ extract was dried over anhydrous sodium sulphate, and the solvent was then distilled out. The residue was dissolved in HPLC-grade methanol before undergoing aglycone moiety analysis. Silver oxide was used to neutralise the acidic aqueous mother liquor, and the precipitate that resulted was filtered out and washed three times with distilled water. The combined filtrate and washings were concentrated using a rotary evaporator under reduced pressure at 50°C. Pyridine and acetic anhydride were used to further acetylate the residue, and GC grade EtOAc was used to prepare the acetylated sample for GC/MS analysis. By comparing the sugar units’ retention times to those of the reference sugar, the sugar units from the GC analysis were verified (acylated). The presence of 32.705 (*β*-D-glucose) and 26.202 (α-L-rhamnose) was confirmed by GC/MS analysis, and additional sugar units were discovered on the bases of NMR data [20].

### 2.5 Biology

#### 2.5.1 Cell culture and growth conditions

Different human cancer cell lines, such as lung (A-549; HOP-62), breast (MCF-7; MDA-MB 231), pancreatic (MiaPaCa-2), colon (SW-620; HCT-116), prostate (PC-3), and neuroblastoma (SH-SY5Y) were grown in growth medium (RPMI-1640 and DMEM) boosted with 10% fetal bovine serum (FBS Qualified: Standard origin Brazil 10270106), streptomycin (100 units/ml) and penicillin (100 units/ml) in tissue culture flasks. Cells were grown in a CO_2_ incubator (Thermocon Electron Corporation, USA) at 37^°^C with permissible atmospheric conditions of (95%) air and (5%) CO_2_ with (98%) humidity. These cell lines were procured from (NCI) National Cancer Institute, USA, and the positive control used in this study was Camptothecin which was obtained from Sigma-Aldrich (Bengaluru, India).

#### 2.5.2 Sulphorhodoamine B (SRB) Assay

The SRB assay is a colorimetric assay performed to evaluate the cytotoxic potential of active inhibitors [21]. 96-well flat transparent plates were seeded with the optimal cell density per well (Flat bottom). 100µL/well of cell suspension of each cell line was plated with their specific cell number, such as PC-3 (7000), MCF-7 (8000), MDA-MB 231(7500), A-549 (7500), MiaPaCa-2 (6500), SW-620 (7500) HCT-116 (7000), HOP-62 (7500) and allowed to grow overnight at 37^°^Cwith 5% Co_2_ in a cell-culture incubator. The cells were treated with different concentrations of test compounds (1, 5, 10, 30, and 50µM) after 24 h of incubation, along with Camptothecin as a positive control. Again, the cells were incubated at similar culture conditions for 48 h, and ice-cold TCA fixed the cells for 1 h at 4^°^C, plates were washed thrice with water and allowed to dry. Further, at room temperature, 0.4% SRB dye was added (100 µl in each well) for half an hour. The plate was then washed thrice with water and once with 1% v/v acetic acid to remove the unbound SRB and allowed to dry for some time at room temperature. In order to solubilise the bound dye100µl of 10mM Tris buffer (pH – 10.4) was added to each well. Plates were then kept on shaker for 5 minutes to dissolve the protein-bound dye properly. Finally, the OD was noted at 540nm in a microplate reader, and then IC_50_ was calculated using GraphPad Prism Software.

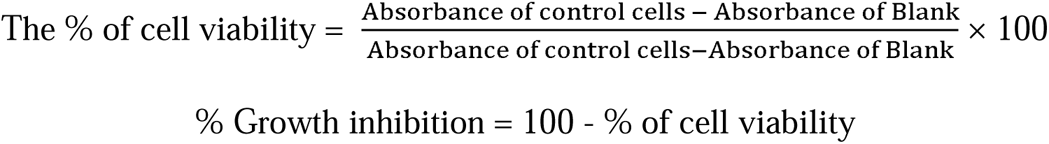

#### 2.5.3 DAPI Staining

The A-549 cells were seeded in 6 well plates (1×10^5)^ for 24 h and were treated with different concentrations of Compound (**2**) for 48 h. After 48 h, PBS wash was given, cell fixation was done using cold methanol, and plates were kept at 4^°^C for 20 minutes. Further, cells were stained with DAPI 1µg/mL in PBS and the cells were observed under a fluorescence microscope (Olympus IX53) to access the morphological changes in nucleus.

#### 2.5.4 Reactive oxygen species (ROS) assay

6 well plates were used for the culture of A-549 cells (2 ×10^5^) for 24 h. After 24 h of incubation, different concentrations of compound (**2**) for 48 h were given. 0.05% H_2_O_2_ was used as a positive control. Post PBS wash, cells staining was done using DCFH-DA (Dichlorodihydro-fluorescein diacetate) dye and were analysed using a fluorescence microscope (Olympus IX53)

#### 2.5.5 Colony formation assay

Cell seeding of A-549(2×10^5^) was done. After 24 h of incubation cells were treated with Compound (**2**) with concentrations of 1, 2, 4, and 6 µM. Trypsinization of cells was done after 48 h and cells were re-seeded as 1000 cells/well. To test the clonogenic potential cells were allowed to regrow to form colonies. Fixation of cells was done with 4% formaldehyde (1 mL/well) followed by PBS wash. Staining was done with 0.5% crystal violet dye and colonies were counted [22].

#### 2.5.6 In vitro cell migration assay

A-549 cells were seeded in 6 well-flat transparent plates and allowed to confluent up to 70-80% for 24 h in starving condition (Serum Starved) and a straight horizontal line with a sterile 200µl tip was created by scrapping the monolayer of A-549 cells. Finally, cells were treated with Compound (2), at concentration of 1, 2, 4, and 6 µM for 48 h. Images of the wounded area were taken at 0 h and 48 h, and the following equation demonstrated the percentage of wound

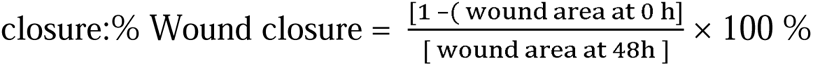

#### 2.5.7 Mitochondrial membrane potential (MMP)

6 well plates were used for the culture of A-549 cells (1.5 ×10^5^) for 24 h and were treated with Compound (2) at 1, 2, 4, and 6 µM and 0.25µM Camptothecin for 48 h. One hour before from terminating the experiment, 10nM Rhodamine −123 dye was added, and PBS wash was given. Cells were analysed under a fluorescence microscope (Olympus IX53) [23].

#### 2.5.8 Western blot analysis

For western blotting, A-549 cells were seeded at a density of 3 × 10^3^cells/ well in 6-wellplates. After 24h, A-549 cells were treated with different concentrations of Compound (2). The concentrations were 1, 2, 4, and 6 µM for 48 h. After 48 h, cells were washed with cold PBS. Cell lysates were prepared in 1X RIPA buffer (Sigma) with added sodium orthovanadate (100mM), protease cocktail (Roche), NaF (100mM), PMSF (100mM) and EDTA (100mM). Protein estimation was done using Bradford reagent (Bio-Rad). The samples were boiled with sample buffer containing 1% *β*-mercaptoethanol, 6%glycerol, 2% SDS, 22mM Tris-HCl pH-6.8, and bromophenol blue. Whole-cell lysates corresponding to 70-100 ug of protein were loaded. SDS-PAGE analysed the samples with a 10% separating gel [24]. After running the gel, the proteins were transferred to the PVDF membrane and then blocked for 1 h in a solution of 5% BSA, 0.1% Tween 20, 150mM NaCl, and 20mM Tris-HCl pH-7.4. After blocking, the membrane was probed with primary BAX, BCL-2, and Cleaved Caspase-3 which was further incubated with corresponding (CST)HRP conjugated secondary antibodies. *β*-Actin was used as a gel loading control.

#### 2.5.9 Statistical analysis

Data analysis was done using MS-Excel and GraphPad Prism5 Software. All data analysis was done using GraphPad Prism-5 Software and Image-J. The statistical significance of data was determined by one-way analysis of variance (ANOVA) and was accepted at P<0.05.

## 3 Results and Discussion

### 3.1 Chemistry

Crude chloroform and 20% aq. MeOH extracts of the rhizomes of *Trillium govanianum*wereimperilled to a sequenceof column chromatographic purification steps using silica gel, HP-20, HP-20SS resins and Sephadex respectively, to obtain two new steroidal saponins, named trilliumoside A (1) and B (2) along with 7 known previously reported compounds.Comparison of their NMR and MS data with the reported literature confirmed the structures of 7 known compounds as Diosgenin [16], Govanic acid [11], 20-Hydroxyecdysone [11], 5,20-Hydroxyecdysone [11], Borassoside E [16,19], GovanosideB [17] and Protodioscin [18,25].Moreover, characterization data of both new compounds is described under.

**Compound 1(TG-07 B3)** was isolated as a brownish solid. Its molecular formula C_51_H_85_O_23_was deduced by LC/MS data at (*m/z* 1064.13 [M+H]^+^ (calcd for C_51_H_84_O_23_, 1063.1911), along with its ^1^H and ^13^C NMR data (Tables 1 and 2). The NMR spectra of compound 1(TG-07B3) exhibited two typical angular signals for two quaternary methyl’s resonating at *δ*_H_0.86 and 1.03 assignable to the (*s*, CH_3_-18 and *s*, CH_3_-19 methyl groups) one methyl doublet for secondary methyl group at *δ*_H_0.92 (*d, J*, 6.3Hz, 21-CH_3_), four oxymethylene protons at *δ*_H_3.41(*m*, H-26a), *δ*_H_3.72(*m*, H-26b), *δ*_H_3.37(*m*, H-27a) and *δ*_H_ 3.70(*m*, H-27b), and their ^13^C respective signals were found resonating at *δ*_C_ 58.43 and 76.12, one olefinic proton resonating at *δ*_H_ 5.38 (*br., s*, H-6), in ^13^C there are two olefinic signals observed at *δ*_C_ 141.95 (C-5) and 122.75 (C-6) and onequaternarycarbon characteristic of hemiketalicfunctuionality found resonating at *δ*_C_112.46 (C-22). The ^1^H NMR spectrum of compound **1** also revealed the presence of four characteristic anomeric proton signals resonating at *δ*_H_ 4.23 (*d, J*-7.Hz, *O*-Glc C-26), 4.49 (*d,*7.84 Hz, *O*-Glc1-H-1), 4.85 (*brs*, Rha-1) and 5.18 (*brd*, *J*-8.7 Hz, Rha-2) while the corresponding anomeric carbon signalswere observed at *δ*_C_ 104.66, 100.5, 103.04 and 102.5 in ^13^C NMR spectra as assigned by HSQC spectra. The ^13^C NMR spectrum exhibited a total of 51 carbon resonances where 27 signals correspond to aglycone part and the remaining 24 carbon resonances were assigned to the sugar units having four six membered monosaccharide units. Moreover, the comparison of the NMR of compound **1** with that of Trikam steroids showed that they share the identical spirostanol skeleton of (*25R*)-5en-spirost-3□,17*α*,27-triol,which was further confirmed by closer examination of 2D NMR data consisting of HSQC, HMBC, COSY and NOESY spectra of compound1 [19].The site of attachment of different sugar units was determined by their key HMBC correlations (Figure 1). The key HMBC correlation of Glc-H-1 (*δ*_H_ 4.49) with C3 (*δ*_C_ 79.45) of aglycone confirmed Glc1’ to be attached at C-3 of the aglycone moiety. There is another HMBC correlations observed between C-4’of Glc1’ (*δ*_H_ 4.49) and Rha-H-1’’ (*δ*_H_ 4.85) established Rha1’’ unit to be attached at C-4 of Glc1’. Further the close examination of HMBC spectrum showed correlation between C-3 of Rha-H-1’’ (*δ*_C_ 79.32) with H-1’’’of Rha-2 (*δ*_H_5.18) confirming that the site of attachment of Rha2 to be at C-3 of Rha1. There is another correlation of O-Glc-H’’’ (*δ*_H_ 4.23) and C-26 (*δ*_C_ 76.11). In addition to NMR data the presence of sugar units was further confirmed by acid hydrolysis of compound **1** which resulted in liberation of two units each of *D*-glucose and *L*-rhamnose. It was done by relating their^1^H and^13^C NMR data and coupling constants with the literature reports. Based on above data, the structure of compound **1** was assigned as 27-O-*β*-D-glucopyranosyl-(25S)-5-en-spirost-3*β*,17α,27-triol-3-*O*-α-L-rhamnopyranosyl–(1-4)-[α-*L*-rhamnopyranosyl-(1-4)]-*β*-*D*-glucopyranoside (TrilliumosideA).

**Figure 1:**
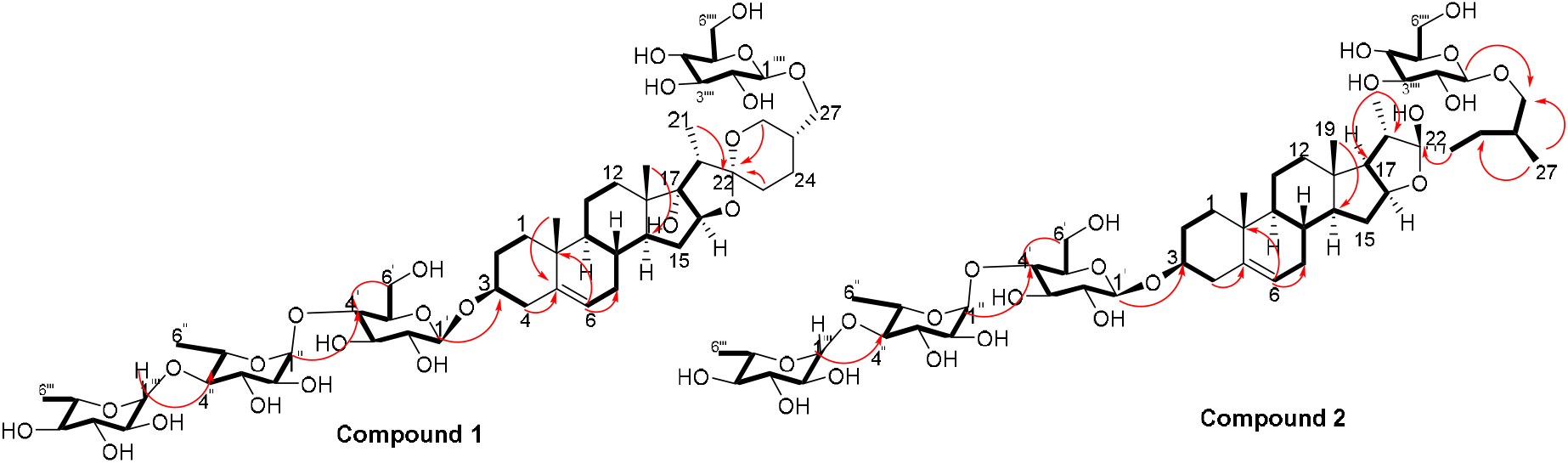
The important HMBC 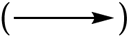 and ^1^H-^1^H COSY 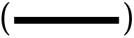 correlations of 1 and 2

**Compound 2 (TG-09)** was obtained as a yellow brownish solid. Its molecular formula was determined to be C_51_H_85_O_22_, by HR-ESI-MS data at (*m/z* 1049.5532 [M+H]^+^ (calcd for C_51_H_84_O_22_, 1048.5411), in combination with ^1^H and ^13^C NMR data (Tables 1 and 2). The analysis of ^1^H and ^13^C NMR spectral dataof compound **2**revealedthe presence of two typical angular signals for two quaternary methyl’s resonating at *δ*_H_0.82 and 1.04 assignable to the (s, CH_3_-18 and s, CH_3_-19 methyl groups, two methyl doublets for two secondary methyl groups at *δ*_H_0.96 (*d, J*, 6.2 *Hz*, CH_3_-27) and 1.20 (*d, J,*6.3*Hz*, 21-CH_3_). There is also characteristic signal for two oxymethylene protons observed at *δ*_H_3.32(m, H-26a) and *δ*_H_ 3.73(m, H-26b) and an olefinic proton resonating at *δ*_H_ 5.38 (*br, s*, H-6). In ^13^C NMR data two olefinic signals were found resonating at *δ*_C_ 141.95 (C-5) and 122.75 (C-6) while there is one characteristic quaternary hemiketalic carbon at *δ*_C_114.46 (C-22), further it was confirmed that the configuration of C-22 hydroxyl group is (□) by its resonance at 114.46 instead of (*α*) at 112 [19]. The configuration 25*R* was confirmed by the chemical shift difference between the geminal protons H-26a (*δ*3.38, m) and H-26b (*δ* 3.70, m) (Δ*δ*ab = 0.32 ppm): Δ*δ*ab<0.48 ppm for 25*R* and Δ*δ*ab> 0.57 ppm corresponds for 25*S* [26]. The ^1^H NMR spectrum of compound 2 also showed signals characteristic for anomeric protons resonating at *δ*_H_4.24 (*d, J-*7.8 Hz, Glc C-26), 4.49 (*d, J*-7.84*Hz*, Glc1-H-1), 4.85 (*brs,* Rha1) and 5.18 (*brd, J-*8.7 *Hz*, Rha 2) with the corresponding anomeric carbon resonances observed at *δ*_C_104.66, 100.5, 103.04 and 102.5 respectively, in ^13^CNMR spectrum. The sugarcomponentwas found to be a composed of two units of *D*-glucose and *L*-rhamnose,obtainedafter acid hydrolysis of compound 2. It was also concluded by comparing their ^1^H and^13^C NMR data and coupling constants with literature reports. The sugar attachment siteswere determined by key HMBC correlations (Figure 2). The presenceofHMBC correlation between Glc-H-1’ (*δ*_H_ 4.49) with C3(*δ*_C_ 79.92) of aglycone proved the Glc1’ to be attached at C-3 of the aglycone. There was another key HMBC correlation observed between C-4’of Glc1’ (*δ*_H_ 4.49) and Rha-H-1’’ (*δ*_H_ 4.85) which confirmedthatRha1 to be located at C-4 of Glc1’. Further the attachment 1’’’of Rha-2 (*δ*_H_5.18) with C-3 of Rha-H-1’’ (*δ*_C_ 79.43) was confirmed by correlations found in HMBC spectrum. There is another correlation of *O*-Glc-H’’’’(*δ*_H_ 4.24) and C-26 (*δ*_C_ 76.11). After detailed analysis of this spectral data the structure of**2** was elucidated as (25*R*)-furost-5-en-3*β*,22*β*,26 triol-3-*O*-α-*L*-rhamnopyranosyl-[(1-4)-α-L-rhamnopyranosyl-(1-4)]-*β*-*D*-glucopyranosyl 26-*O*-*β*-D-glucopyranoside (Trilliumoside B).

**Figure 2:** Structure of compounds (1-9)

### 3.2 Biology

#### 3.2.1 Cell growth inhibition studies

The anti-cancer potential of crude extracts (CHCl_3,_ 20% aq. MeOH and one sub-fraction of 20% aq. MeOH) and isolated compounds were evaluated for their *in-vitro* cytotoxic potential, which was expressed in percentage of inhibition and IC_50_ values respectively (Table 3 and 4) against a panel of human cancer cell lines namely lung cancer (A-549, HOP-62), breast (MCF-7, MDA-MB 231), pancreatic (MiaPaCa-2), colon cancer (SW-620, HCT-116), prostate (PC-3) and Neuroblastoma (SH-SY5Y) cancer using SRB assay. The results revealed that the extracts and fraction EF (IC_50_µg/mL) exhibited the highest *in-vitro* cytotoxicity, followed by 20%aq. MeOH and CHCl_3_(IC_50_µg/mL). However, among the pure compounds, Compound 1 (IC_50_µM) and Compound 2 (IC_50_µM) showed maximum inhibition effect on A-549 and SW-620 cell lines in a concentration-dependent manner (Table 3). Further, three known compounds, Diosgenin, Protodioscin, and Borassoside E, showed good cytotoxic activity on A-549 and SW-620.However, compounds (TG-01, TG-03, TG-04, TG-05, TG-06, and TG-12) at a concentration of 10µM did not possess significant cytotoxicity (< 45 %) against any of the tested human cell lines. Compounds (1) and (2) showed the best % growth inhibition of the all isolated compounds, 75.35 and 92.11 at 10µM and 50µM.

**Table 3:** Growth inhibitory effect of Extracts and enriched fractions against different cancer cell lines

| Extract & Fractions | Conc. (µg/mL) | Human cancer cell lines from five different tissues |  |  |  |  |  |  |
| --- | --- | --- | --- | --- | --- | --- | --- | --- |
|  |  | Breast | Breast | Colon | Colon | Lung | Pancreatic | Prostate |
|  |  | MCF-7 | MDAMB-231 | HCT-116 | SW-620 | A-549 | MIA PaCa-2 | PC-3 |
|  |  | Growth Inhibition (%) |  |  |  |  |  |  |
| **TG-E1** | **100** | 16.88 | 28.55 | 11.83 | 10.97 | 12.49 | **ND** | **ND** |
|  | **50** | 5.24 | 11.87 | 8.39 | 3.22 | 4.56 | **ND** | **ND** |
| **TG-E3** | **100** | 48.22 | 40.28 | 72.91 | **72.50** | **66.46** | 30.87 | 67.75 |
|  | **50** | 23.07 | 18.76 | 50.73 | 58.87 | 35.07 | 15.74 | 52.26 |
| **TG-EF** | **100** | 60.22 | 58.22 | 86.66 | **91.29** | **96.30** | 68.87 | 85.99 |
|  | **50** | 45.40 | 34.78 | 76.77 | 75.39 | 82.78 | 42.48 | 66.55 |

**Table 4:** SRB assay-based screening results. **IC_50_ values (µM) of compounds on panel of different human cancer cell lines.

| IC<sub>50</sub> (μM)± SD |  |  |  |  |  |  |  |  |  |  |
| --- | --- | --- | --- | --- | --- | --- | --- | --- | --- | --- |
| Compd | A-549 | HCT-116 | MCF-7 | MDA-MB 231 | MiaPaCa-2 | SW-620 | PC-3 | SHSY-5Y | HOP-62 | FR2 |
| 1 | 1.83±0.54 | 2.91±0.50 | 4.40±0.33 | 1.90±0.26 | 1.94±0.24 | 1.85±0.52 | 3.18±0.37 | 2.43±0.32 | 1.92±0.62 | 13.75±0.42 |
| 2 | 1.79±0.47 | 3.47±0.39 | 5.92±0.55 | 9.49±0.25 | - | 3.18±0.40 | 4.97±0.58 | - | - | 10.84±0.35 |
| 3 | >50 | >50 | >50 | >50 | >50 | >50 | >50 | >50 | >50 | - |
| 4 | 2.08±0.56 | 1.95±0.45 | 6.98±0.39 | 8.29±0.33 | 3.79±0.49 | 1.98±0.46 | 3.77±0.52 | 3.33±0.26 | 2.65±0.47 | - |
| 5 | 1.50±0.62 | 4.44±0.57 | 4.15±0.47 | 2.56±0.33 | 1.69±0.45 | 1.66±0.38 | 1.85±0.43 | 4.43±0.44 | 1.85±0.57 | 2.71±0.23 |
| 6 | >50 | >50 | >50 | >50 | >50 | >50 | >50 | >50 | >50 | - |
| 7 | 30.46±0.49 | >50 | 29.40±0.37 | >50 | >50 | >50 | >50 | >50 | >50 | - |
| 8 | >50 | >50 | >50 | >50 | >50 | >50 | >50 | >50 | >50 | - |
| 9 | 32.95±0.43 | >50 | 38.99±0.63 | >50 | 33.26±0.57 | >50 | >50 | >50 | >50 | - |
| CPT | 0.030±0.04 | 0.160±0.23 | 0.147±0.13 | 0.125±0.03 | 0.060±0.02 | 0.093±0.03 | 0.170±0.03 | - | 0.065±0.04 | - |

Out of both the bioactive compounds (1) & (2), compound (2) was further evaluated for anticancer studies by performing DAPI, ROS, MMP, Colony formation assay, and Wound healing / Scratch assay. <u>But due to the restricted quantity of compound (2), we could not proceed</u> <u>with its in vivo studies. However, compound (1) was produced in adequate quantity and carried</u> <u>forward for in</u> vitro and *invivo* studies and the results for the same will be published in the near future.

#### 3.2.2 Compound (2) altered nuclear morphology assessed by DAPI staining

Apoptotic cells have characteristic features like nuclear shrinkage, chromatin condensation and formation of apoptotic bodies [27]. DAPI (4’,6-diamidino-2-phenylindole) [28–29] dye was used to assess the nuclear changes. This dye is specific to minor grooves of the A-T region of DNA which was analysed by using fluorescence microscopy. In this experiment, treatment of Compound (2) on A-549 was given at 1, 2, 4, and 6 µM concentrationsfor 48 h. Camptothecin (CPT)was used as a positive control. Apoptotic bodies and chromatin condensation was seen in the nuclei of A-549 cells treated with Compound (2)ina concentration dependent manner. These characteristic features were absent in untreated control cells (Figure 4). These features clearly depicts the anti-proliferative effect of compound (2)

**Figure 3:**
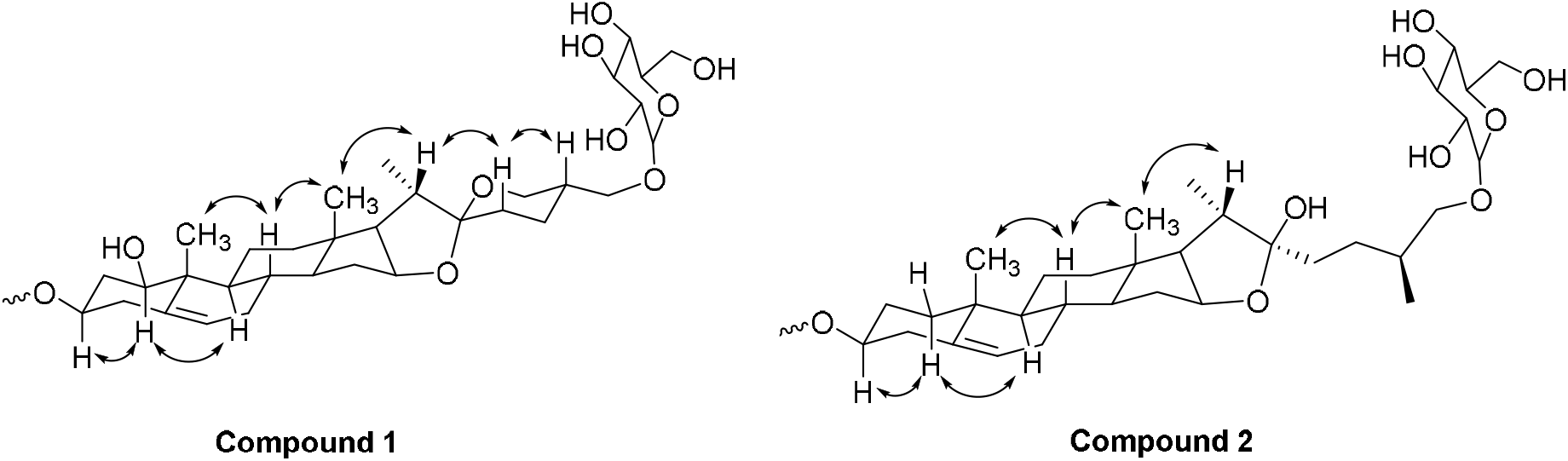
The key^1^H-^1^H NOESY (arrow) correlations of **1** and **2**

**Figure 4:**
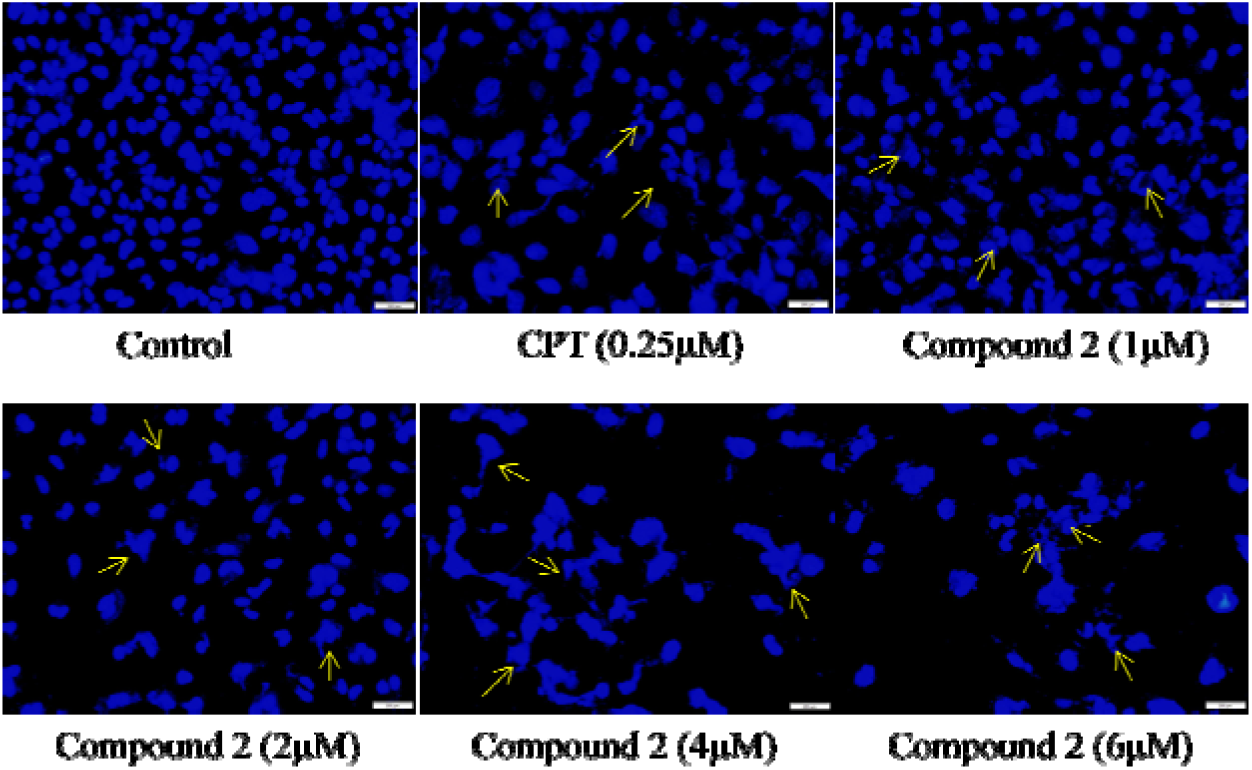
DAPI assay in A-549 cells observed under fluorescence microscope to determine nuclear morphological changes.

#### 3.2.3 Intracellular ROS production induced by Compound (2) in A-549 cells

The presence of high quantities of reactive oxygen species (ROS) is a hallmark of programmed cancer cell death. Increased intracellular ROS level damage the membrane lipids, intra-cellular proteins, organelles, and nucleic acids, which ultimately causes cell death [30]. In this study, A-549 cells were seeded at 2×10^5^ per ml/well density. After 24 h, cells were treated with Compound (2) at the concentrations of 1, 2, 4, and 6 μM. Later, staining of cells was done with DCFDA (dichlorodihydro-fluorescein diacetate) dye.Fluorescence microscopy was used toanalyze the reactive oxygen species ROS. A sharp increase in fluorescence intensity wa observedcompound treated as well as in positive control (Hydrogen peroxide (H_2_O_2_). Presence of ROS was confirmed by fluorescent product dichlorofluorescein (DCF) in dose dependent manner.

#### 3.2.4 Compound (2) lowers mitochondrial membrane potential (MMP)

Loss of mitochondrial membrane permeability and release ofcytoplasmic apoptogenic stimuli lead to cell death [31]. A fluorescent dye, rhodamine-123, was used to analyse the change in MMP. An Olympus IX53 imaging microscope monitored the quenching of fluorescence. The loss of mitochondrial membrane integrity is directly related to the fluorescence decay rate. Mitochondrial membranedestabilization causes the leakage of rhodamine-123, which lowers the intensity of fluorescence [32]. The MMP levels of A-549 cells are significantly decreased after treatment with compound 2 at 1, 2, 4, and 6 μM concentrations. Compound (2) inducesthe depolarization of mitochondrial membrane potential(low ΔΨmt) in concentration dependent manner.

#### 3.2.5 Compound (2) inhibited in vitro cell migration during wound healing assay in A-549 cells

In this experiment, A-549 cells monolayer were scratched and treated with different concentrations of Compound (2), i.e., 1, 2, 4, and 6 µM for 48 h. Images were taken at 0 and 48 h in order to quantitatively assess the percentage reduction in cell migration [33]. It was observed that inhibition of cell migration took place in concentration-dependent manner as compared to control. This indicates the potential of Compound (2) to retard cancer cell growth.

#### 3.2.6 Compound (2) inhibited cell proliferation during colony formation assay in lung cancer cells (A-549)

Cells’ proliferation capabilities (the potential of single cell to grow into a colony) were measured by *in vitro* clonogenic assay [34]. A-549 cells were seeded in a 6-well flat transparent well and treated with Compound (2) of various concentrations of 1, 2, 4, and 6 µM. Compound (2) significantly inhibited colony formation in A-549 cells as the concentration increases along with inhibition effect (Figure 8).

**Figure 5:**
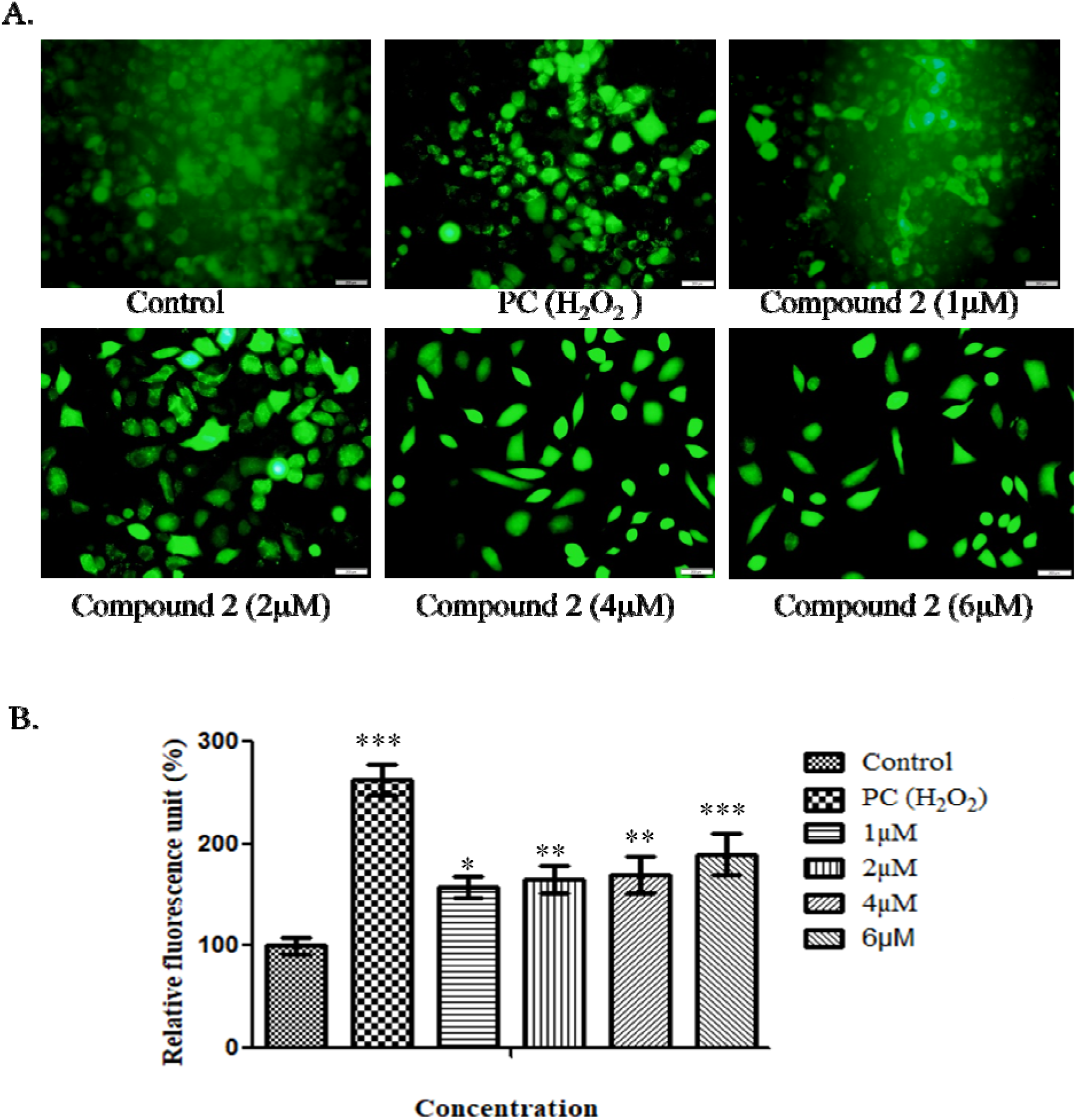
(A)ROS production in A-549 cells following treatment with different concentrations of Compound (2). As a positive control, H_2_O_2_ is used. In turn, concentration enhances the fluorescence of ROS production. (B) Bar graph of ROS production.Data was calculated from 3 independent experiments with p<0.05.

**Figure 6:**
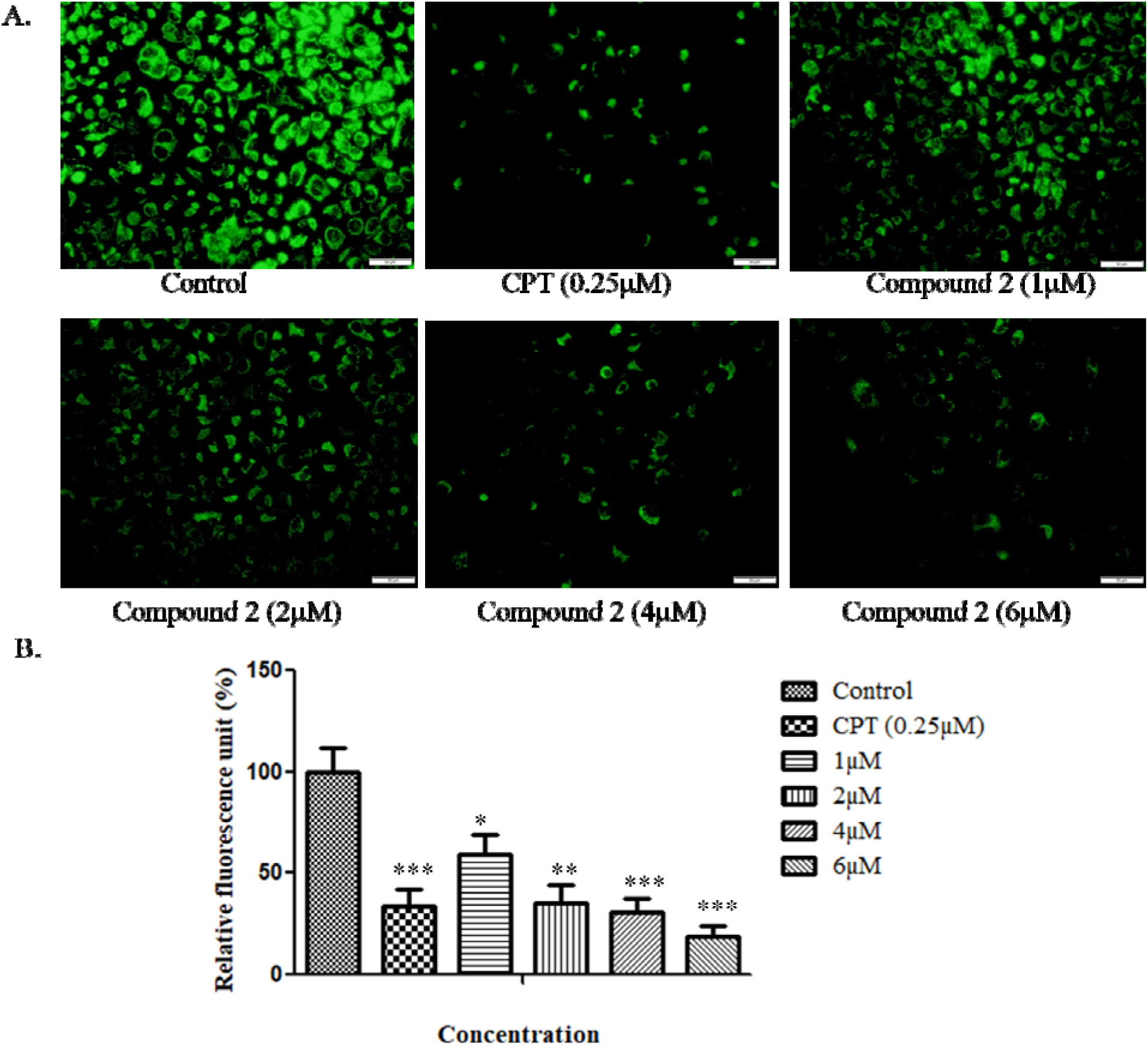
(A)Dose dependent loss of MMP in A-549 cells treated with Compound 2 for 48 h analyzed through fluorescence microscopy. Camptothecin was taken as a positive control. (B) Bar graph of ROS production.Data was calculated from 3 independent experiments with p<0.05.

**Figure 7:**
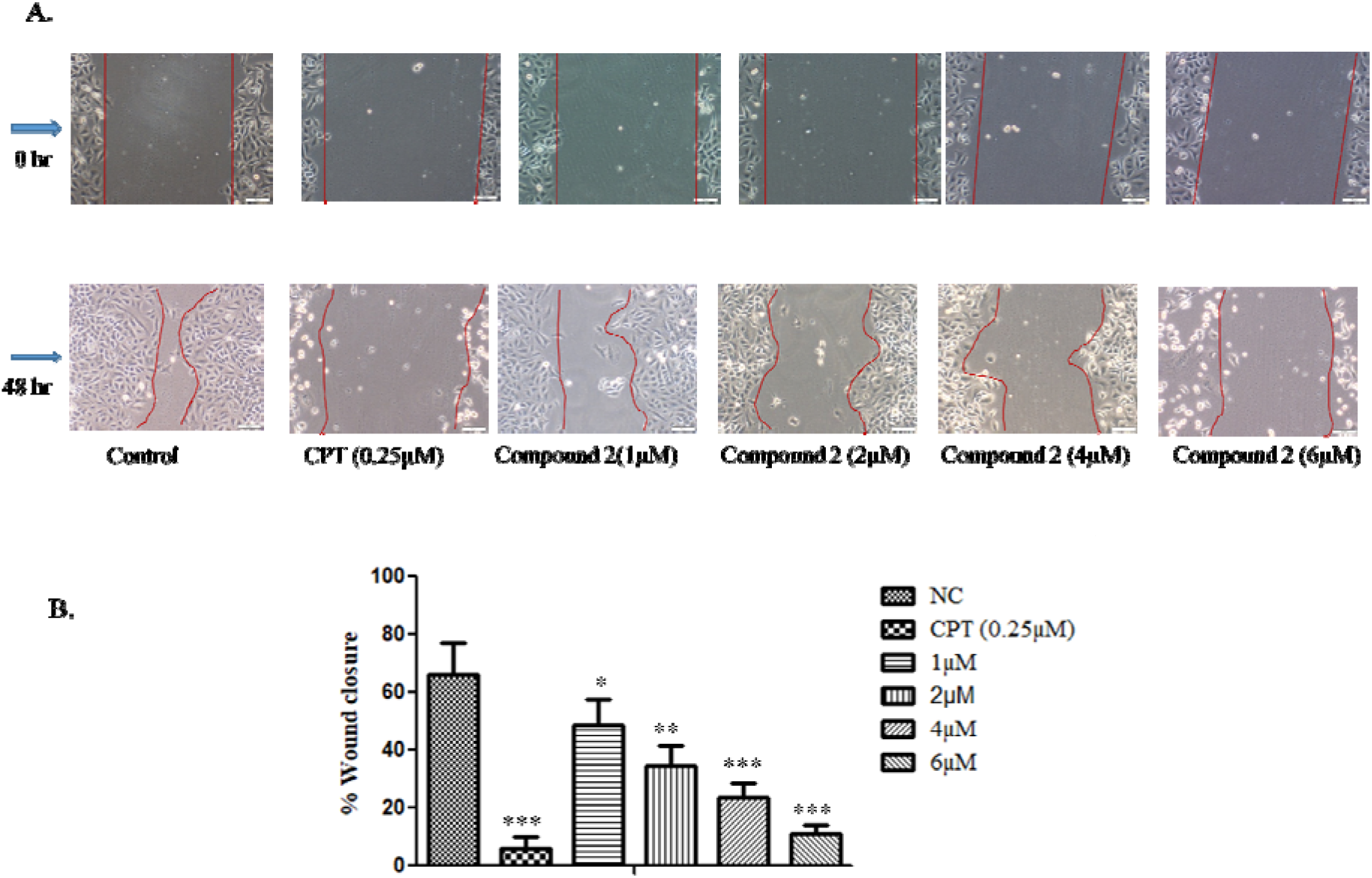
*In vitro* wound healing assay- (A) A-549 cells treated with Compound 2 for 48 h reduces the migration of the cell as the concentration of compound was increased (B) Wound closure area with control using ImageJ software. Data was calculated from 3 independent experiments with p<0.05.

**Figure 8:**
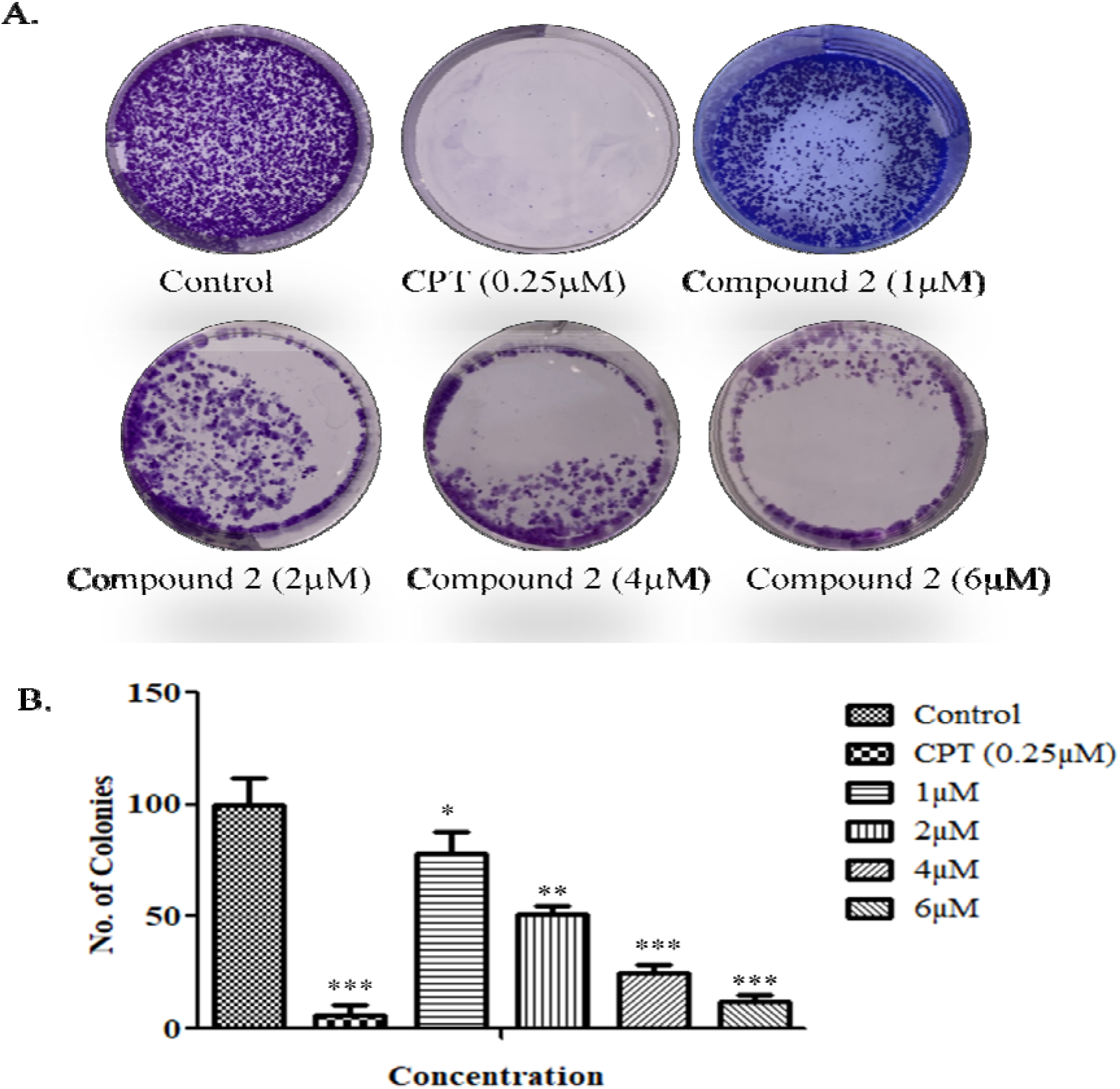
(A)Colony assay was performed over A-549 cells. (B) Number of colonies was counted and plotted as a bar graph. Camptothecin (0.25µM) was used as Positive control. Data was calculated from 3 independent experiments with p<0.05.

#### 3.2.7 Western blotting

For the verification of apoptosis induction by Compound (2) in A-549 cells, western blotting was performed with BCL-2, BAX, and Cleaved Caspase 3. BCL-2 which is a pro-proliferative marker, showed consistent downregulation in dose-dependent manner[35]. On the other hand, steady increase has been observed in the expression of pro-apoptotic proteins viz., BAX and Caspase 3(Figure 9).

**Figure 9:**
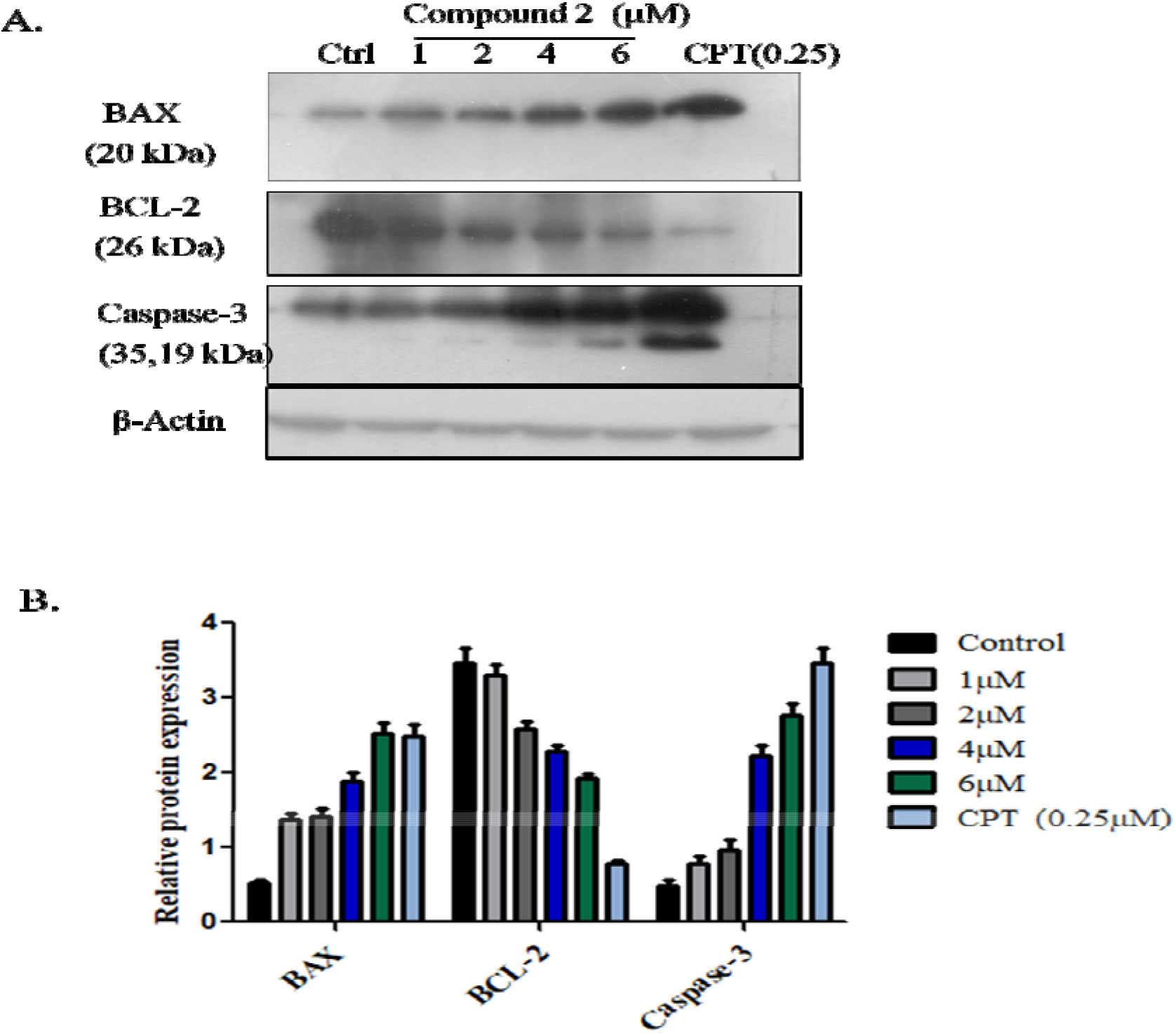
(A) Immunoblotdepiction BAX, BCL-2 and cleaved Caspase-3 in A-549 cells by Compound 2 for 48 h(B) represents the dentiometric analysis of western blots. Data was calculated from 3 independent experiments with p<0.05.

## 4. Conclusion

Over the past years, *Trillium govanianum* has gained the attention of researchers because of very less exploration of its phytochemistry and pharmacological studies, especially in the area of cancer research. In our present study, we isolated and characterized two new steroidal saponins from the rhizomes of *Trillium govanianum.* Compounds (1) and (2) exhibited significant cytotoxic activity against human lung and colon cancer cell lines in a considerable micromolar range. Trilliumoside A (1) showed significant cytotoxicity with IC_50_ values of 1.83 and 1.85 µM on A-549 (Lung) and SW-620 (Colon) cell lines, whereas Trilliumoside B (2) IC_50_ value against A-549 cell line was found to be 1.79 µM. Mechanistic anticancer assays were performed on compound (2) which revealed noteworthy changes like MMP reduction, increase in ROS production, inhibition of anti-apoptotic protein BCL-2, and activation of BAX and Caspase-3. All these changes were responsible for the induction of apoptosis in the A-549 cancer cell line which will lay the foundation of compound (2) as a lead molecule in anticancer drug discovery.

## 5. DATA AVAILABILITY STATEMENT

The original contributions presented in the study are included in the article/Supporting Information; further inquiries can be directed to the corresponding author.

## 6. AUTHOR CONTRIBUTIONS

BL carried out isolation, characterization of compounds and wrote original draft.MT designed and performed *in-vitro* anti-cancer activity. AB helped in characterization of compounds.UD helped in manuscript preparation. DM and PN supervised the pharmacological experiments. SG provided and the identified the plant material. PG conceptualized, supervised, coordinated all the studies and finalised the manuscript. All the authors read and approved the final manuscript.

## 8. ACKNOWLEDGMENTS

Authors are thankful to the Director CSIR-IIIM for his interest in this research work. BL (ICMR), MT (UGC), AB (CSIR), UD (CSIR), PG, DM, PN (CSIR) thankfully acknowledge the financial support in the form of research fellowships. The manuscript bears institutional communication number CSIR-IIIM/IPR

## 9. SUPPORTINGINFORMATION

The Supporting Informationfor this article can be found online at: Supporting InformationSpectroscopic data of compounds 1 and 2, including 1D (^1^Hand ^13^C), 2D (^1^H-^1^H COSY, HSQC, HMBC, and NOESY), NMR and ESIMS.

