## Supplementary file for "Trilliumosides A-B, two novel steroidal saponins isolated from Rhizomes of *Trillium govanianum* as potent anticancer agents targeting apoptosis in A-549 cancer cell line"

**Bashir Ahmad Lone^a,b^, MisbahTabassum^a,c^, Anil Bhushan^a,b^, Urvashi Dhiman^a,b^, Prem N. Gupta^a,c^, D.M. Mondhe^a,c^,SumeetGairola^a,d^ ,PrasoonGupta^a,b*^**

^a^Academy of Scientific and Innovative Research, CSIR-Human Resource Development Centre, Campus Ghaziabad, 201002, India

^b^Natural Products and Medicinal Chemistry Division CSIR-Indian Institute of Integrative Medicine, Canal Road Jammu -180001, J&K, India

^c^Pharmacology Division, CSIR-Indian Institute of Integrative Medicine, Jammu 180001, India

^d^Plant Science and Agrotechnology Division CSIR-Indian Institute of Integrative Medicine, Canal Road Jammu -180001, J&K, India

*Corresponding author (s):

Dr. Prasoon Gupta, Principal Scientist, Natural Products and Medicinal Chemistry Division

CSIR-Indian Institute of Integrative Medicine, Canal Road Jammu -180001, J&K, India

**Abstract**

Two new steroidal saponins, Trilliumoside A (1) and B (2) were isolated by bioactivity-guided phytochemical investigation from 20% aqueous-methanol extract from the rhizomes of *Trillium govanianum* along with seven previously known compounds such as protodioscin (3) govanoside B (4), borassoside E (5), 20-hydroxyecdysone (6), 5-20-hydroxyecdysone (7), govanic acid (8), and diosgenin (9). The structure elucidation of new compounds was carried out using spectroscopic methods such as 1D, 2D NMR data and HR-ESI-MS. The isolated compounds were evaluated for *in-vitro* cytotoxic activity against a panel of different human cancer cell lines. Compound (1) showed significant cytotoxicity with IC_50_ values of 1.83 and 1.85 µM on A-549 (Lung) and SW-620 (Colon) cell lines, whereas compound (2) IC_50_ value against A-549 cell line was found to be 1.79 µM. Among previously known compounds (3), (5) and (9) their cytotoxic IC_50_ value was found to be in the range of 5-10 µM. In detailed anticancer analysis compound (2) was seen inhibiting colony forming potential and invitro migration in the A-549 cell line. Furthermore, the mechanistic study of compound (2) on the A-549 cell line revealed characteristic changes including nuclear morphology, increased ROS generation, and reduced levels of MMP. Above mentioned events eventually induce apoptosis, a key hallmark in cancer studies, by upregulating the pro-apoptotic protein BAX and downregulating the anti-apoptotic protein BCL-2 thereby activating Caspase-3. Our study reports the first mechanistic anticancer evaluation of the compounds isolated from the rhizomes of *Trillium govanianum* with remarkable activity in the desired micromolar range.

**Figure S1.** ^1^HNMR spectrum of compound **1(TG-07 B3)**(MeOD_6_, 500 MHz)

**Figure S2.**^13^C NMR spectrum of compound **1** (MeOD_6_, 100 MHz)

**Figure S3.** DEPT spectrum of compound **1**(MeOD_6_, 100 MHz)

**Figure S4.** HSQC spectrum of compound **1**

**Figure S5.** HMBC spectrum of compound **1**

**Figure S6.** COSY spectrum of compound **1**

**Figure S7.** NOESY spectrum of compound **1**

**Figure S8.** LCMS spectrum of compound **1**

**Figure S9.** ^1^HNMR spectrum of compound **2 (TG-09)**(MeOD_6_, 500 MHz)

**Figure S10.** ^13^C NMR spectrum of compound **2** (MeOD_6_, 500 MHz)

**Figure S11.** DEPT spectrum of compound **2**(MeOD_6_, 500 MHz)

**Figure S12.** HSQC spectrum of compound **2**

**Figure S13.** HMBC spectrum of compound **2**

**Figure S14.** COSY spectrum of compound **2**

**Figure S15.** NOESY spectrum of compound **2**

**Figure S16.** HR-ESIMS spectrum of compound **2**

Compound-1(TG-07 B3)

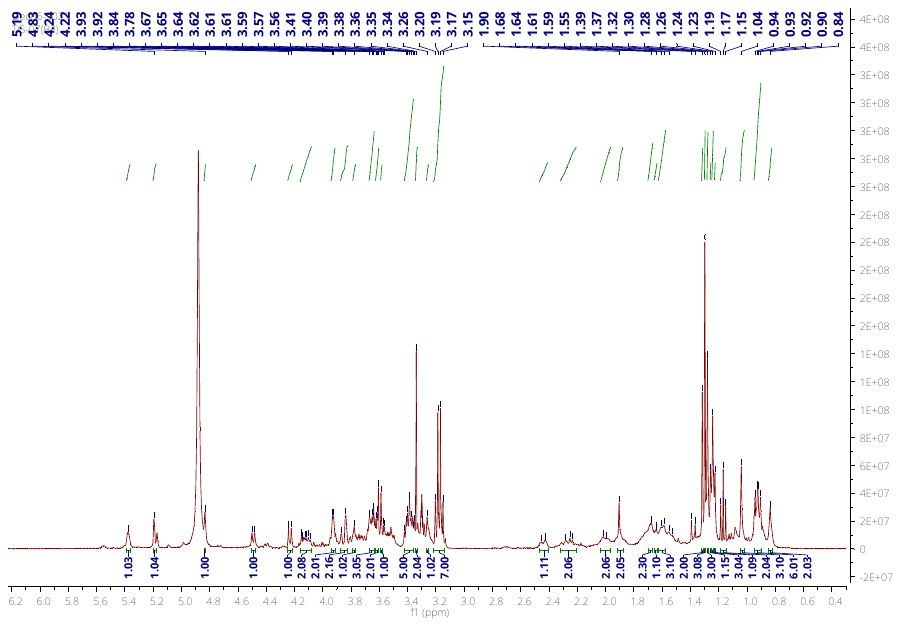

**Figure S1.** ^1^H NMR (400 MHz, CD_3_OD) of compound TG 07 B3

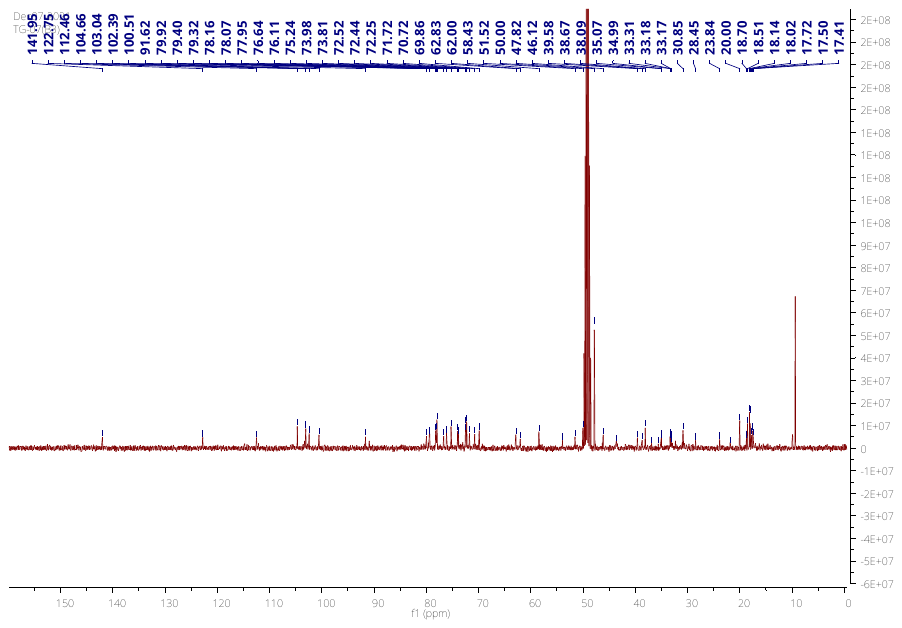

**Figure S2.** ^13^C NMR (400 MHz, CD_3_OD) of compound TG 07 B3

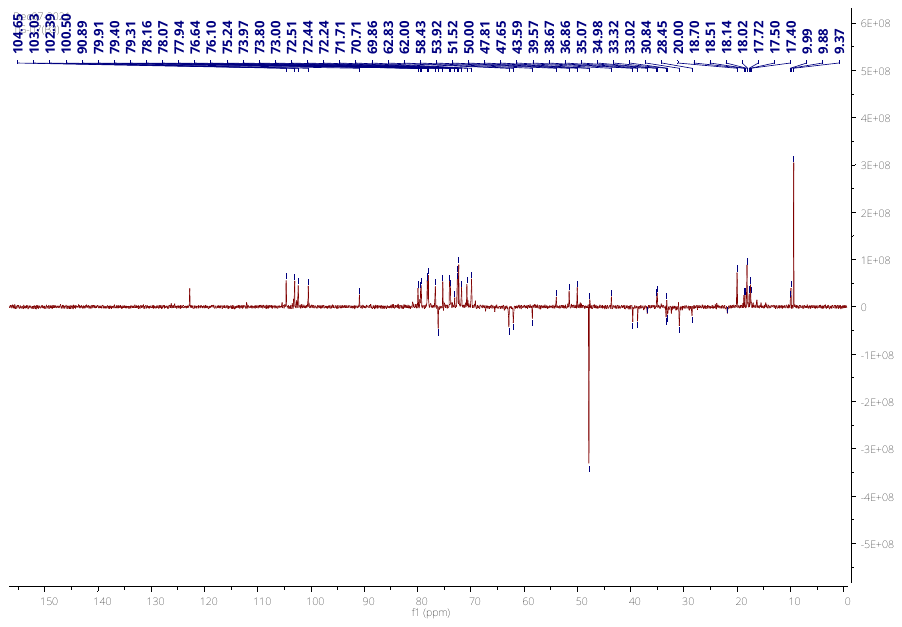

**Figure S3.** DEPT Spectra of compound TG 07 B3

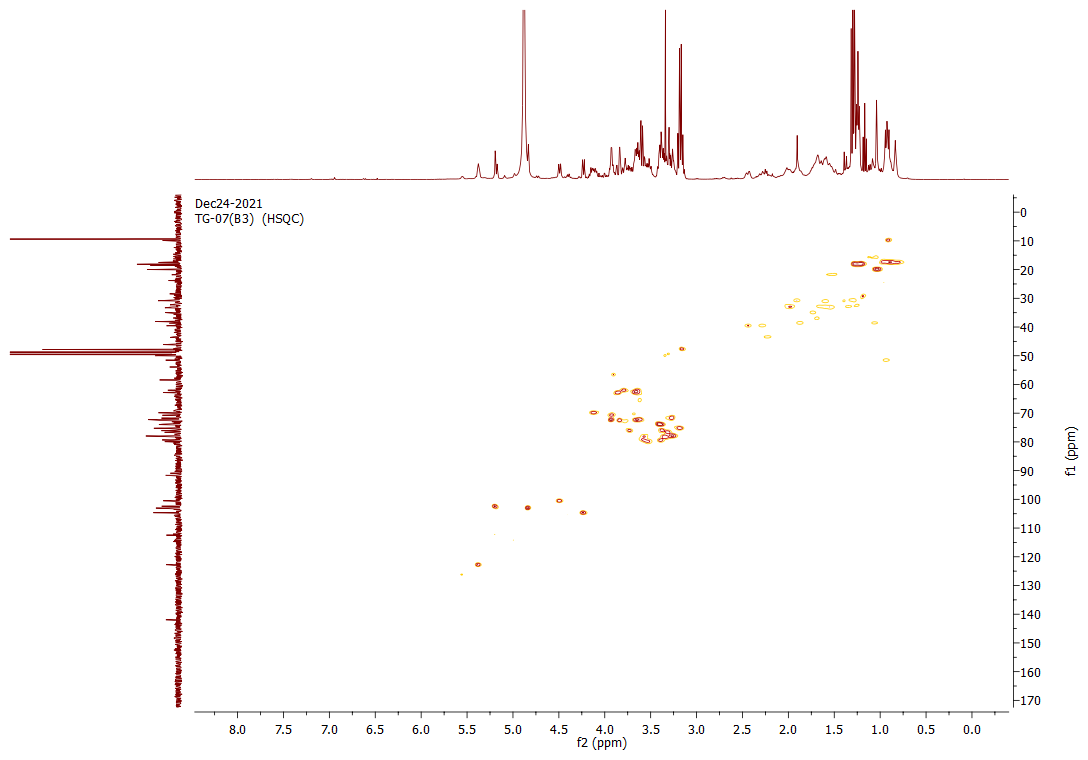

**Figure S4.** HSQC Spectra of compound TG 07 B3

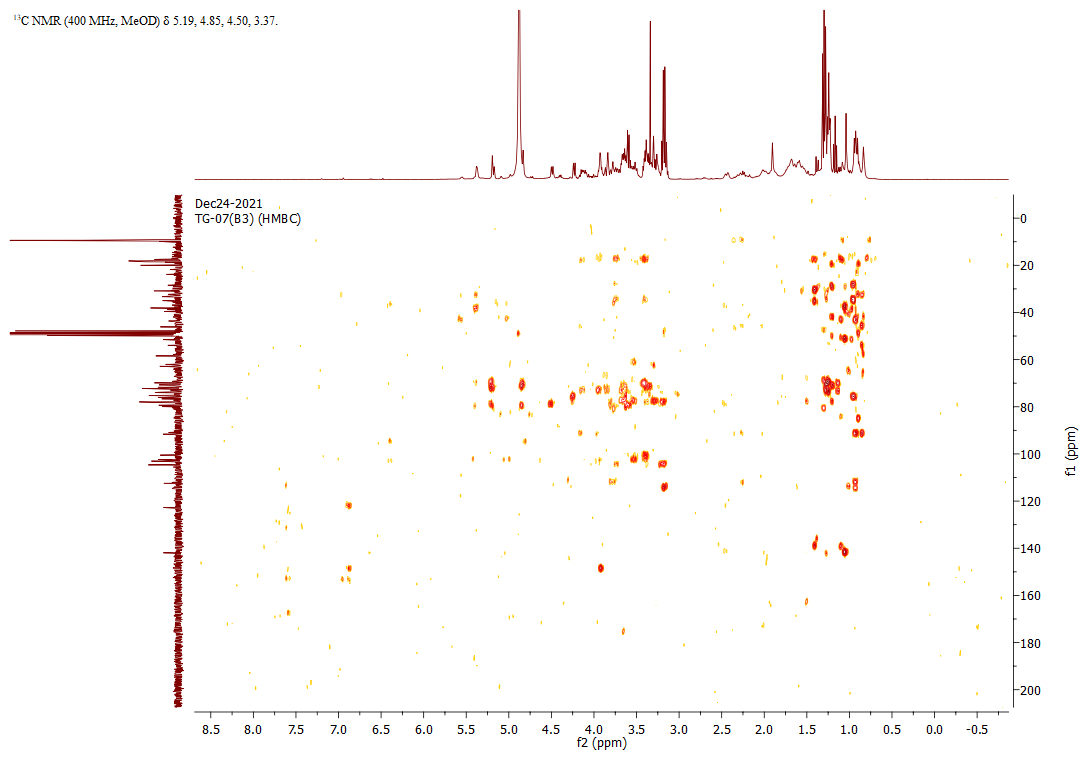

**Figure S5.** HMBC Spectra of compound TG 07 B3

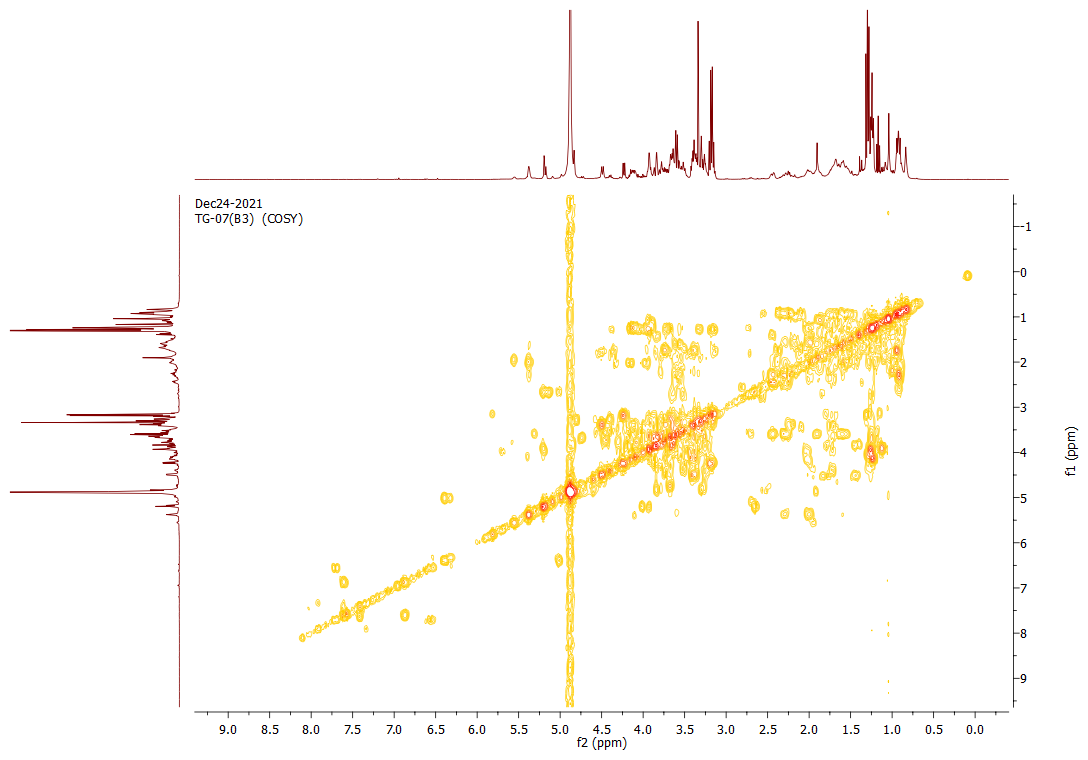

**Figure S6.** COSY Spectra of compound TG 07 B3

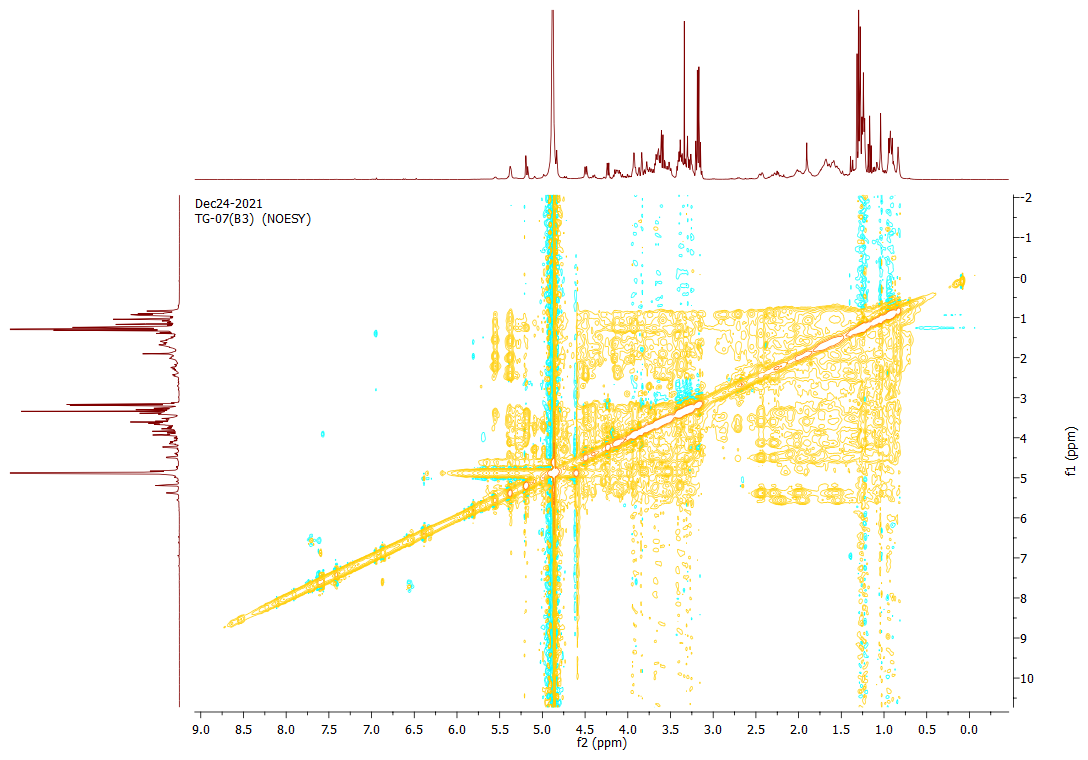

**Figure S7.** NOESY Spectra of compound TG 07 B3

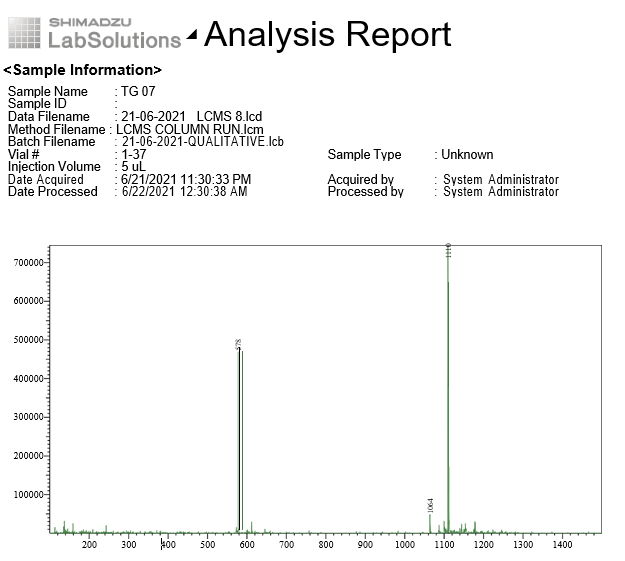

**Figure S8.** LC/MS of Compound 1

**Compound-2(TG-09)**

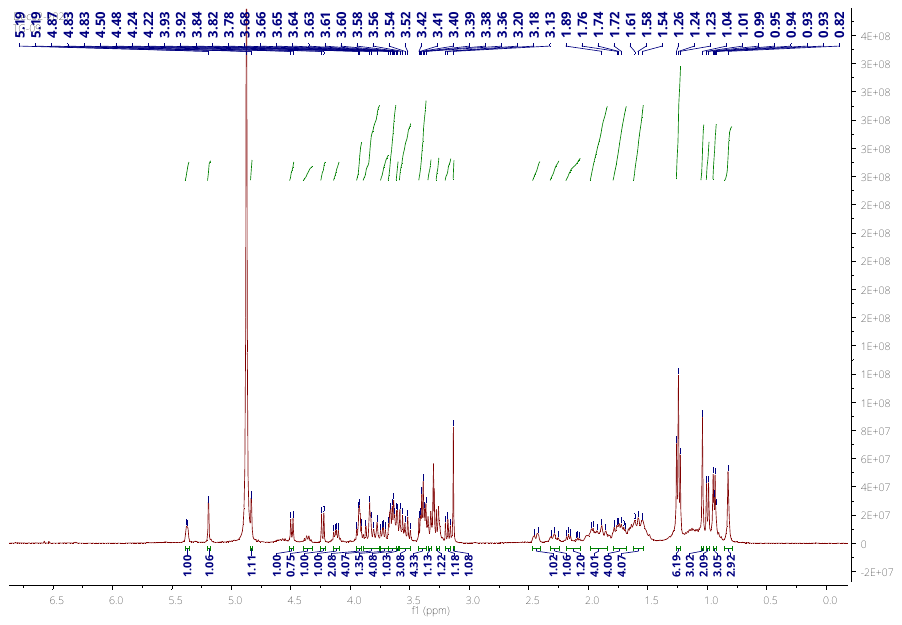

**Figure S9.** ^1^H NMR (400 MHz, CD_3_OD) of compound TG 09

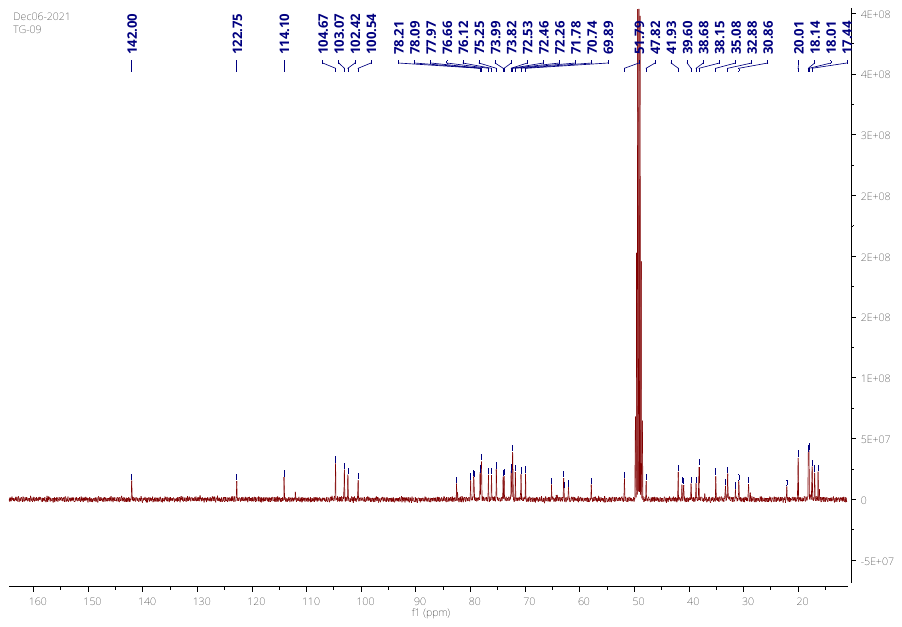

**Figure S10.** ^13^C NMR (400 MHz, CD_3_OD) of compound TG 09

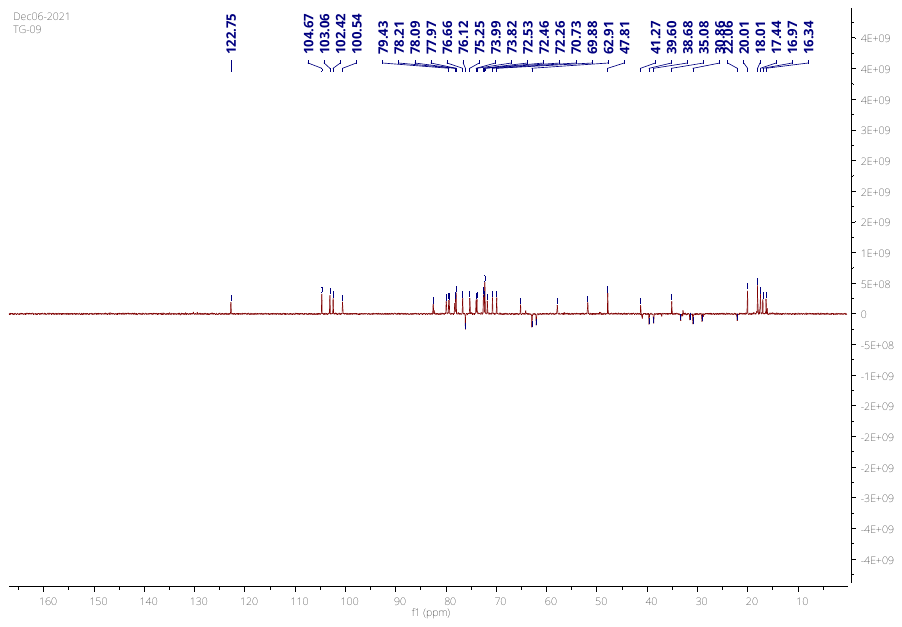

**Figure S11.** DEPT Spectra of compound TG 09

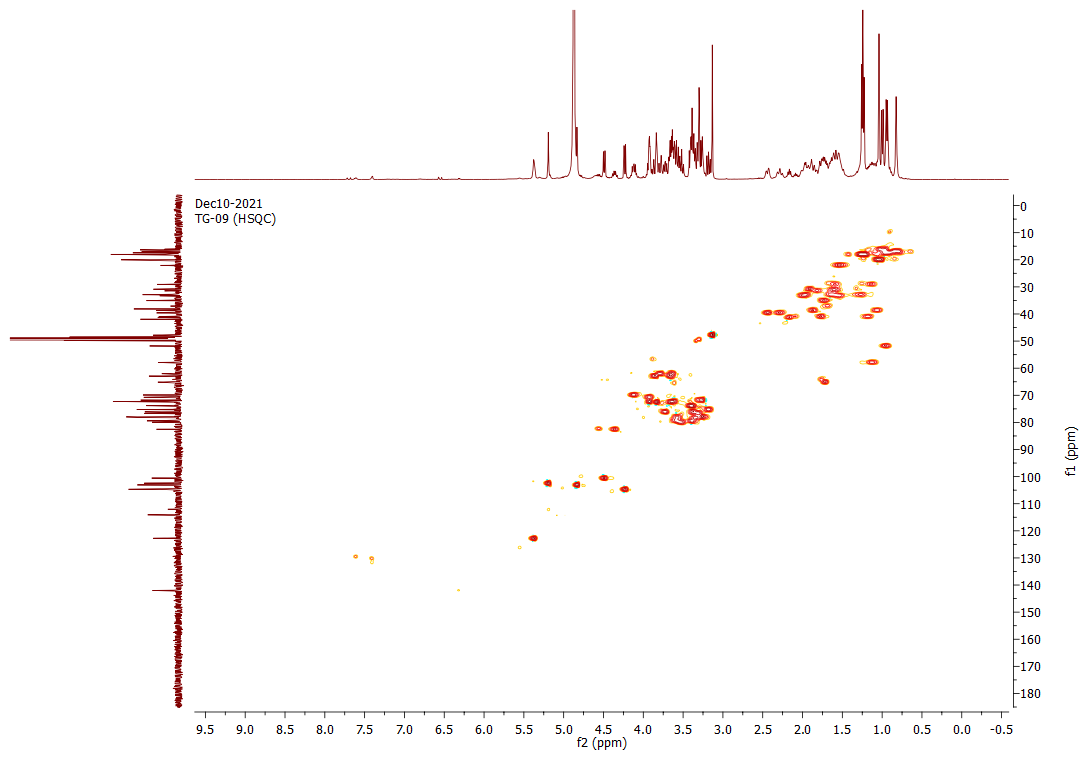

**Figure S12.** HSQC Spectra of compound TG 09

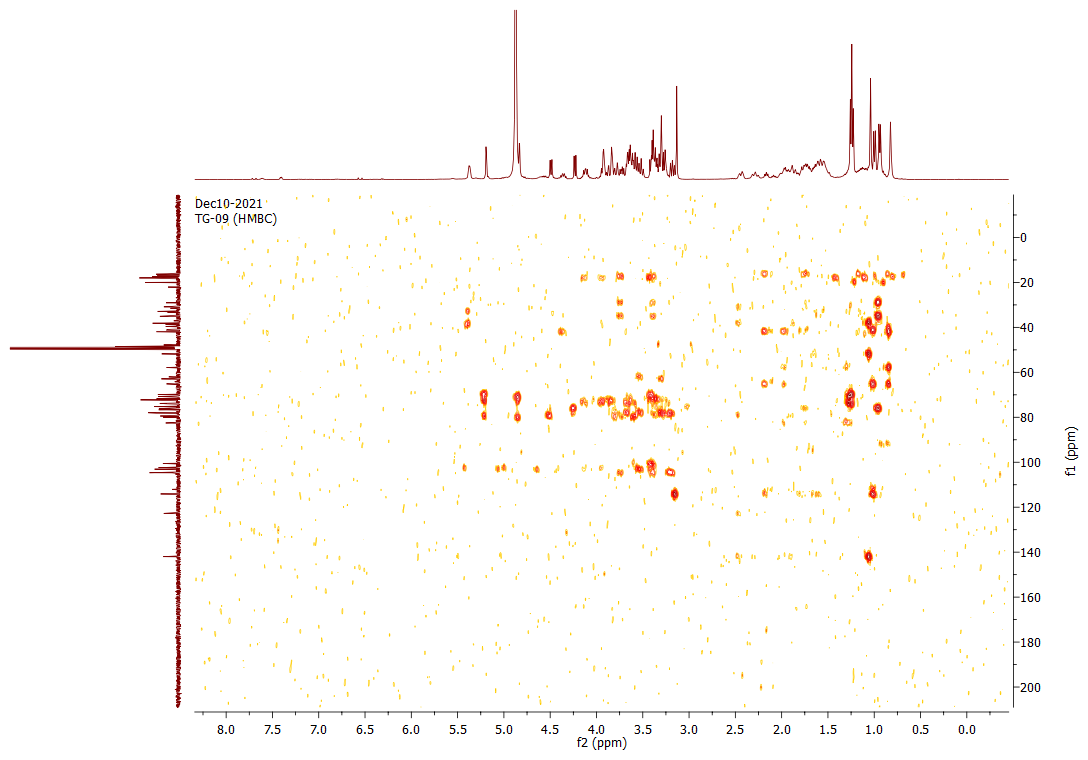

**Figure S13.** HMBC Spectra of compound TG 09

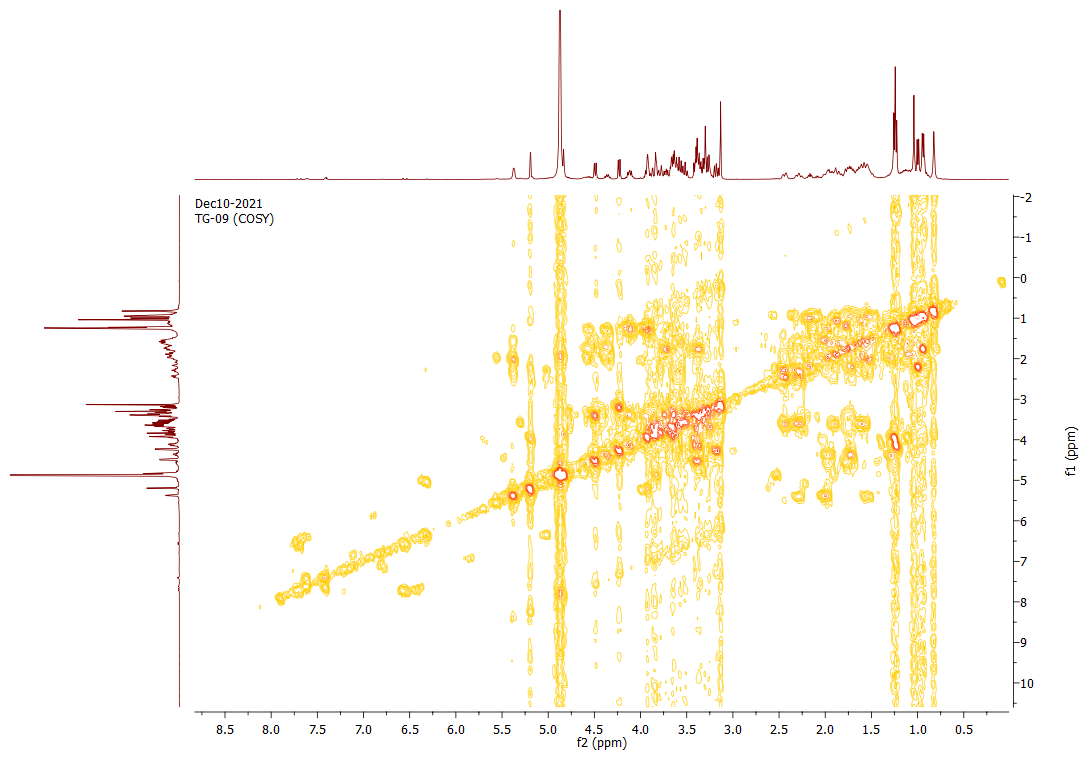

**Figure S14.** COSY Spectra of compound TG 09

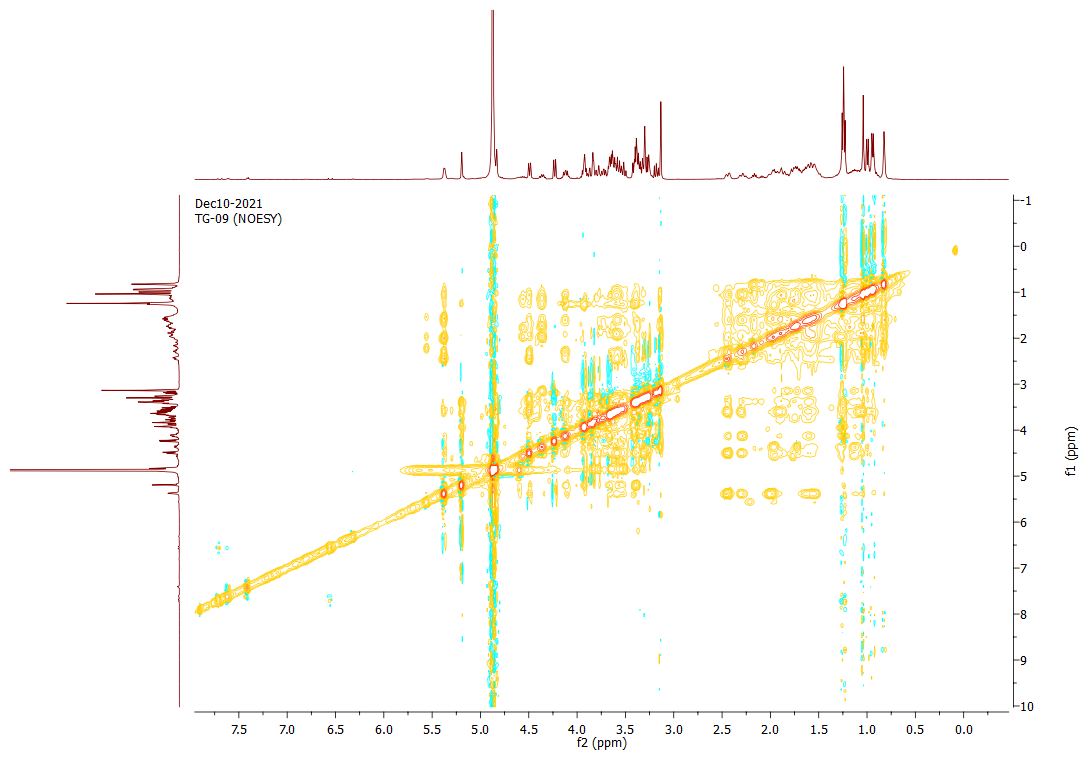

**Figure S15.** NOESY Spectra of compound TG 09

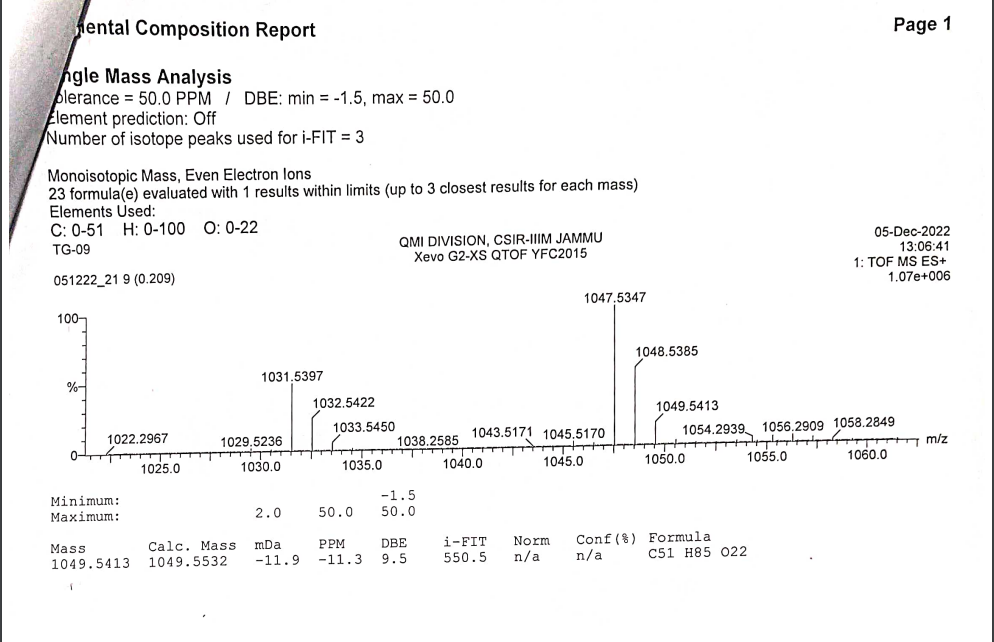

**Figure S16.** HR-ESIMS of compound TG 09
